# Xrs2 and Tel1 independently contribute to MR-mediated DNA tethering and replisome stability

**DOI:** 10.1101/425942

**Authors:** Julyun Oh, So Jung Lee, Rodney Rothstein, Lorraine S. Symington

**Affiliations:** Biological Sciences Program, Columbia University, New York, New York 10027; Department of Microbiology & Immunology, Columbia University Irving Medical Center, New York, NY 10032; Department of Genetics & Development, Columbia University Irving Medical Center, New York, NY 10032

**Keywords:** Mre11, Rad50, Xrs2, Tel1, DNA repair, DNA replication, genome stability

## Abstract

The yeast Mre11-Rad50-Xrs2 (MRX) complex has structural, signaling and catalytic functions in the cellular response to DNA damage. Xrs2, the eukaryotic-specific component of the complex, is required for nuclear import of Mre11 and Rad50, and to recruit the Tel1 kinase to damage sites. We show that nuclear-localized MR complex (Mre11-NLS) catalyzes homology-dependent repair without Xrs2, but MR cannot activate Tel1 and it fails to tether DSBs resulting in sensitivity to genotoxins, replisome instability and increased gross chromosome rearrangements (GCRs). Fusing the Tel1 interaction domain from Xrs2 to Mre11-NLS is sufficient to restore telomere elongation and Tel1 signaling to Xrs2-deficient cells. Furthermore, Tel1 stabilizes Mre11-DNA association, and this stabilization function becomes important for DNA damage resistance in the absence of Xrs2. Enforcing Tel1 recruitment to the nuclear MR complex fully rescues end tethering, stalled replication fork stability and suppresses GCRs, highlighting important roles for Xrs2 and Tel1 to ensure optimal MR activity.

**Highlights:**

- Xrs2 is required for recruitment but not for activation of Tel1 kinase
- Tel1 and Xrs2 function independently to optimize MR activity at DSBs and stalled replication forks
- Stable association of Mre11 at DSBs is required to maintain end-to-end tethering
- MR-mediated DNA tethering promotes replisome stability and genome integrity

## Introduction

The Mre11-Rad50-Xrs2 (MRX) complex plays a central role in the DNA damage response through detection and repair of cytotoxic DNA double-strand breaks (DSBs). Mutation of genes encoding the MRX complex in *Saccharomyces cerevisiae* causes genotoxin sensitivity, shortening of telomeres and meiotic defects (Borde, 2007; Gobbini et al., 2016). Hypomorphic mutations of human MRN complex components (Nbs1 replaces Xrs2 in mammalian cells) result in the chromosome instability syndromes, Nijmegen breakage syndrome (NBS), NBS-like disease (NBSLD) and ataxia telangiectasia-like disease (ATLD), which are associated with radio-sensitivity, cancer predisposition and immune deficiencies (Carney et al., 1998; Stewart et al., 1999; Waltes et al., 2009). The cellular phenotype of NBS and ATLD are similar to A-T, which is caused by loss of the ATM kinase (Shiloh and Ziv, 2013). In contrast to yeast, the mammalian MRN complex is essential for cell viability (Stracker and Petrini, 2011).

MRX/N is rapidly recruited to DSBs where it tethers DNA ends and activates the Tel1/ATM kinase to signal the DNA damage checkpoint (Stracker and Petrini, 2011). In addition to the signaling role in response to DSBs, recruitment and activation of Tel1 by MRX at telomeres is necessary for telomere elongation (Ritchie and Petes, 2000). Moreover, the complex participates in both of the major DSB repair mechanisms: non-homologous end joining (NHEJ) and homologous recombination (HR). For NHEJ, MRX recruits factors necessary for direct re-ligation of the broken ends (Chen et al., 2001; Matsuzaki et al., 2008; Palmbos et al., 2008). For HR, the complex utilizes its nuclease activity to catalyze degradation of the 5’-terminated strands of the break ends to yield single stranded DNA (ssDNA), substrate for the Rad51 recombinase (Symington et al., 2014). MRX/N associates with unperturbed replication forks and stabilizes stalled forks during replication stress (Dungrawala et al., 2015; Mirzoeva and Petrini, 2003; Sirbu et al., 2011; Tittel-Elmer et al., 2009). Loss of MRX results in extreme sensitivity to hydroxyurea (HU), loss of replisome components from stalled replication forks and impaired fork progression in the presence of HU (Seeber et al., 2016; Tittel-Elmer et al., 2009).

Mre11 and Rad50 are conserved in all domains of life, whereas Xrs2/Nbs1 appears to be restricted to eukaryotes (Stracker and Petrini, 2011). Mre11 has ssDNA endonuclease and 3’-5’ dsDNA exonuclease activities in vitro that are important for end resection. Rad50 is a member of the structural maintenance of chromosome (SMC) family of proteins, characterized by ATPase motifs at the N and C termini separated by a long coiled-coil domain (Stracker and Petrini, 2011). The globular DNA binding domain of the Mre11-Rad50 complex is comprised of an Mre11 dimer associated with the ATPase cassettes of a Rad50 dimer. ATP binding and hydrolysis regulate access of Mre11 to DNA, thereby controlling the nuclease activity (Deshpande et al., 2014; Lammens et al., 2011; Lim et al., 2011; Mockel et al., 2012). The extended coiled-coil domains of Rad50, and Zn-mediated dimerization of the hook domains at the apex of the coiled-coils, are thought to be important to tether DNA ends at DSBs and for sister-chromatid interactions (Hohl et al., 2011; Tittel-Elmer et al., 2012; Wiltzius et al., 2005). Maintaining close proximity of DNA ends may promote NHEJ by stimulating ligation (Chen et al., 2001), while bridging sister chromatids at DSBs facilitates the homology search during HR and prevents the damaged chromatid from physically separating from the rest of the chromosome. Consistent with this view, the integrity of the coiled-coil and Rad50 hook domains are crucial to prevent a DSB from becoming a chromosome break (Hohl et al., 2011; Lobachev et al., 2004; Wiltzius et al., 2005).

Xrs2/Nbs1 is the least conserved member of the complex and is associated with eukaryotic-specific functions, such as NHEJ and DNA damage signaling. Xrs2/Nbs1 interacts with phosphorylated Lif1/Xrcc4 and Sae2/CtIP via the FHA domain at the N-terminus (Liang et al., 2015; Lloyd et al., 2009; Matsuzaki et al., 2008; Palmbos et al., 2008; Williams et al., 2009), and with Mre11 and Tel1/ATM through conserved motifs in the C-terminal region of the protein (Falck et al., 2005; Kim et al., 2017; Limbo et al., 2018; Nakada et al., 2003; Tsukamoto et al., 2005; You et al., 2005). Eukaryotic Mre11 homologs have a large loop insertion within the phosphodiesterase domain, referred to as the latching loop, which mediates interaction with Xrs2/Nbs1 (Park et al., 2011; Schiller et al., 2012). Nbs1 binding to the latching loops extends the Mre11 dimer interface and stabilizes the dimeric form, suggesting that Xrs2 has an active role in the architecture of the MR complex in addition to its role in checkpoint signaling (Schiller et al., 2012). Indeed, expression of just a 108 amino acid fragment of murine Nbs1, encompassing the Mre11 interaction domain, is sufficient to sustain cell viability (Kim et al., 2017). Xrs2/Nbs1 is the only component of the complex harboring a nuclear localization signal (NLS) and its interaction with Mre11 is necessary for translocation of Mre11-Rad50 into the nucleus (Carney et al., 1998; Desai-Mehta et al., 2001; Tsukamoto et al., 2005).

Previous studies have shown that fusing Mre11 to an NLS (Mre11-NLS) partially suppresses the slow growth and DNA damage sensitivity of Xrs2-deficient cells by restoring Mre11 nuclease and Sae2-dependent end resection (Oh et al., 2016; Tsukamoto et al., 2005); however, NHEJ and Tel1 activation are not restored, highlighting the role of Xrs2 as an Mre11 chaperone and scaffold protein, recruiting factors necessary for these functions (Oh et al., 2016). The goal of the current study was to determine the role of Xrs2 in Tel1 activation. We found that fusing the Tel1 interaction domain from Xrs2 to Mre11-NLS (Mre11-NLS-TID) is sufficient to restore telomere elongation and Tel1 signaling to Xrs2-deficient cells. The Mre11-NLS-TID fusion proteins improve Mre11 association with DSBs and further suppress the DNA damage sensitivity of *xrs2Δ* cells. The suppression is dependent on Tel1, but partially independent of the kinase activity, suggesting a structural role of Tel1 in DNA repair. Moreover, *MRE11-NLS xrs2Δ* cells exhibit a severe DNA end tethering defect and instability of stalled replication forks, which are again rescued by enforcing Tel1 recruitment to the Mre11 complex. Together, our data suggest a model whereby Xrs2 and Tel1 independently contribute to Mre11 complex stabilization at DSBs and stalled replication forks to promote genome integrity.

## Results

### Enforcing Tel1 recruitment to the Mre11-Rad50 complex

Our previous study showed that *MRE11-NLS xrs2Δ* cells are unable to recruit and activate Tel1 upon DSB formation, and are defective for telomere maintenance (Oh et al., 2016). In addition, Mre11 enrichment at DSBs is reduced compared to wild-type (WT) cells, similar to a *tel1Δ* mutant (Gobbini et al., 2015). Because Tel1 is required for the normal retention of Mre11 at DSBs, we asked if enforcing Tel1 recruitment in the absence of Xrs2 could restore Tel1 signaling and stabilize the MR complex at DSBs.

To address these questions, we fused the Tel1 interacting domain (TID) of Xrs2 to the C-terminus of Mre11-NLS. A previous study showed that the C-terminal 161 amino acids of Xrs2 are necessary and sufficient for Tel1 interaction (Nakada et al., 2003); however, the precise TID within the C-terminal fragment of Xrs2 is not strictly defined. In *Schizosaccharomyces pombe* and *Xenopus laevis* Nbs1, a highly conserved FXF/Y motif preceded by an acidic patch of amino acids was shown to be essential for Tel1^ATM^ binding (You et al., 2005), and a recent study showed that fusing the C-terminal 60 amino acids of *S. pombe* Nbs1 to Mre11 is sufficient to restore Tel1 signaling to Nbs1-deficient cells (Limbo et al., 2018). *S. cerevisiae* Xrs2 has two such motifs, one located 100 amino acids from the C-terminus and another within the C-terminal 15 amino acids. For this reason, we constructed Mre11-NLS-TID fusion proteins with two differing lengths of the Xrs2 C-terminus: 164 amino acids, consisting of both FXF motifs, and 85 amino acids with only the most C-terminal FXF motif (Fig 1A). The *MRE11-NLS-X164* and *MRE11-NLS-X85* constructs were integrated at the *leu2* locus on chromosome III with the *MRE11* promoter and 3’ UTR sequences in a strain with a deletion of the endogenous *MRE11* locus. Expression of the TID fusion proteins is slightly lower than Mre11, similar to Mre11-NLS (Fig S1A). Because a previous study found that a short C-terminal fragment of Xrs2, including the Mre11 binding domain and Tel1 binding domain (residues 630-854), is able to rescue DNA damage sensitivity and partially restore telomere length when expressed in an *xrs2Δ* background (Tsukamoto et al., 2005), we constructed the same fragment and integrated it into the chromosome with the *XRS2* promoter and 3’ UTR sequences (X224) (Fig 1A). Additionally, we constructed a derivative of the X224 fragment fused to a MYC epitope to compare steady state protein levels to full-length Xrs2-MYC; both proteins are expressed at similar levels (Fig. S1B). There are two predicted NLS sequences in Xrs2, a monopartite NLS at residues 350-360 and a bipartite NLS located at the C-terminus (residues 816-849) of the protein (predicted by cNLS Mapper). The fusion proteins and the X224 fragment all contain the predicted bipartite and lack the monopartite NLS. The observation that X224 is able to partially complement *xrs2Δ* demonstrates that the predicted bipartite NLS alone is able to facilitate nuclear localization of the MRX complex (Tsukamoto et al., 2005).

**Figure 1.**
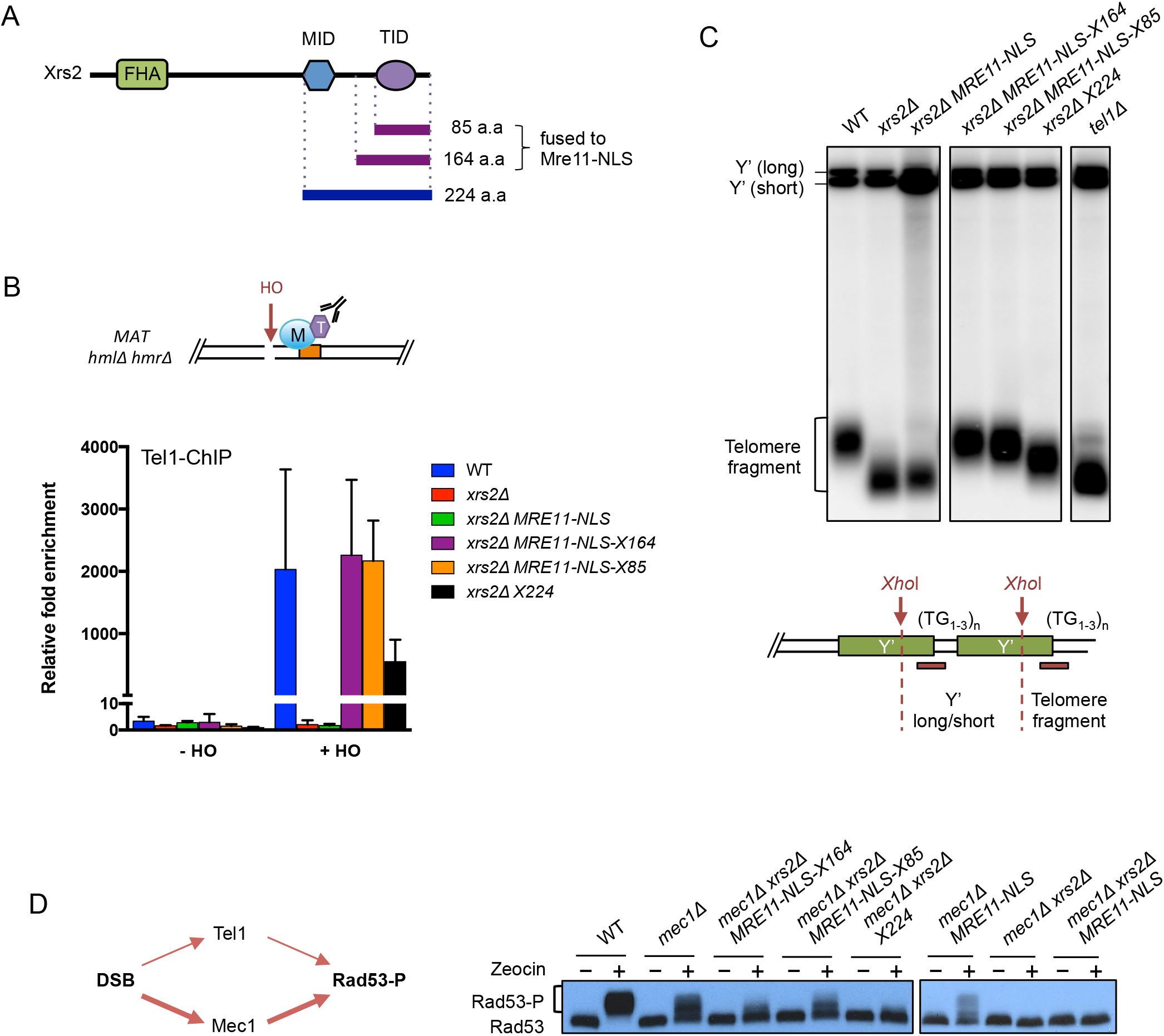
Enforcing recruitment and activation of Tel1 in *xrs2Δ* cells. (A) Schematic representation of Xrs2 protein binding domains and the C-terminal fragments used in this study. FHA: forkhead-associated domain; MBD: Mre11 binding domain; TID: Tel1 binding domain. (B) Schematic representation of the *MAT* locus used in ChIP experiments. The orange bar indicates the region amplified by qPCR. ChIP-qPCR for HA-Tel1 0.2 kb from an HO-induced DSB at the *MAT* locus in cells before (-HO) or 90 min after HO induction (+HO). The error bars indicate SD (n = 3). (C) Southern blot of XhoI-digested genomic DNA hybridized with a Y’ element probe for analysis of telomere lengths. Schematic representation of the telomeric Y’ elements and TG repeats. *XhoI* digestion yield a terminal fragment of ~1.3 kb in WT strains. (D) Model of Rad53 phosphorylation (Rad53-P) in response to DNA damage. Tel1/ATM responds to MRX/N bound DSBs, whereas Mec1/ATR is activated by RPA bound to the ssDNA formed at resected DSBs. Western blot analysis showing Rad53-P in response to 1 hr of zeocin (500µg/ml) treatment.

Recruitment of Tel1 to sequences adjacent to the HO endonuclease cut site at the *MAT* locus was measured by chromatin immunoprecipitation (ChIP). In these strains, the galactose-inducible *GAL1-10* promoter regulates expression of the HO endonuclease, and *HML* and *HMR* are deleted to prevent homology-dependent repair of the DSB. Tel1 binding was measured prior to and 90 min after HO induction. Expression of both of the fusion proteins in the *xrs2Δ* background restores Tel1 enrichment to WT levels, while expression of the *X224* fragment only partially suppresses the *xrs2Δ* Tel1 recruitment defect (Fig 1B). Consistently, telomeres are restored to WT length in cells expressing the fusion proteins while the *X224* fragment only partially rescues the short telomere phenotype (Fig 1C). To examine Tel1 activity in response to DNA damage, phosphorylation of the downstream effector kinase Rad53 was measured following acute zeocin treatment. Rad53 is activated by Tel1 bound to MRX at DSBs or by Mec1-Ddc2 associated with RPA-coated ssDNA generated as a result of end resection. Because the Mec1 pathway is dominant in yeast, it was necessary to use *mec1Δ* strains to detect Rad53 activation by the Tel1 pathway (all the strains also have a *sml1Δ* mutation to suppress lethality caused by *mec1Δ* (Zhao et al., 1998)). Cells expressing the fusion proteins show reduced but visible Rad53 phosphorylation while Rad53 does not show an obvious mobility shift in cells expressing the *X224* fragment (Fig 1D). These data indicate that fusion of TID to Mre11-NLS is able to recruit and activate Tel1 in the absence of Xrs2, and that the *X224* fragment has reduced ability to recruit and activate Tel1 compared to the fusion proteins.

### Tel1 stabilizes Mre11 at DSB ends and enhances DNA damage resistance in the absence of Xrs2

*MRE11-NLS xrs2Δ* and *tel1Δ* strains show decreased retention of Mre11 at DSBs (Cassani et al., 2016; Gobbini et al., 2015; Oh et al., 2016). To address whether recruiting Tel1 to the MR complex could restore enrichment of Mre11, we measured Mre11 binding to sequences adjacent to the HO cut site by ChIP. Consistently, expression of the fusion proteins, as well as the *X224* fragment, in the *xrs2Δ* mutant rescues the defective retention of Mre11 at DSBs (Fig 2A). Surprisingly, expression of all three constructs results in higher Mre11 enrichment than observed in WT cells. We speculated that the higher level of Mre11 is due to a role for Xrs2 in turnover of the complex. Since the FHA domain is missing in all three constructs, we measured Mre11 enrichment in the *xrs2-SH* mutant, which contains mutations of two conserved residues within the FHA domain. Indeed, the *xrs2-SH* mutant shows a similar increased enrichment of Mre11 to the fusion proteins and the *X224* fragment, suggesting that the FHA domain of Xrs2 plays a role in eviction of the MRX complex from DSB ends (Fig 2A). Deletion of *TEL1* in the *xrs2Δ X224* strain completely abolishes the restoration of Mre11 retention, indicating that Tel1 is responsible for the observed increased enrichment of Mre11 at DSBs (Fig 2A).

**Figure 2.**
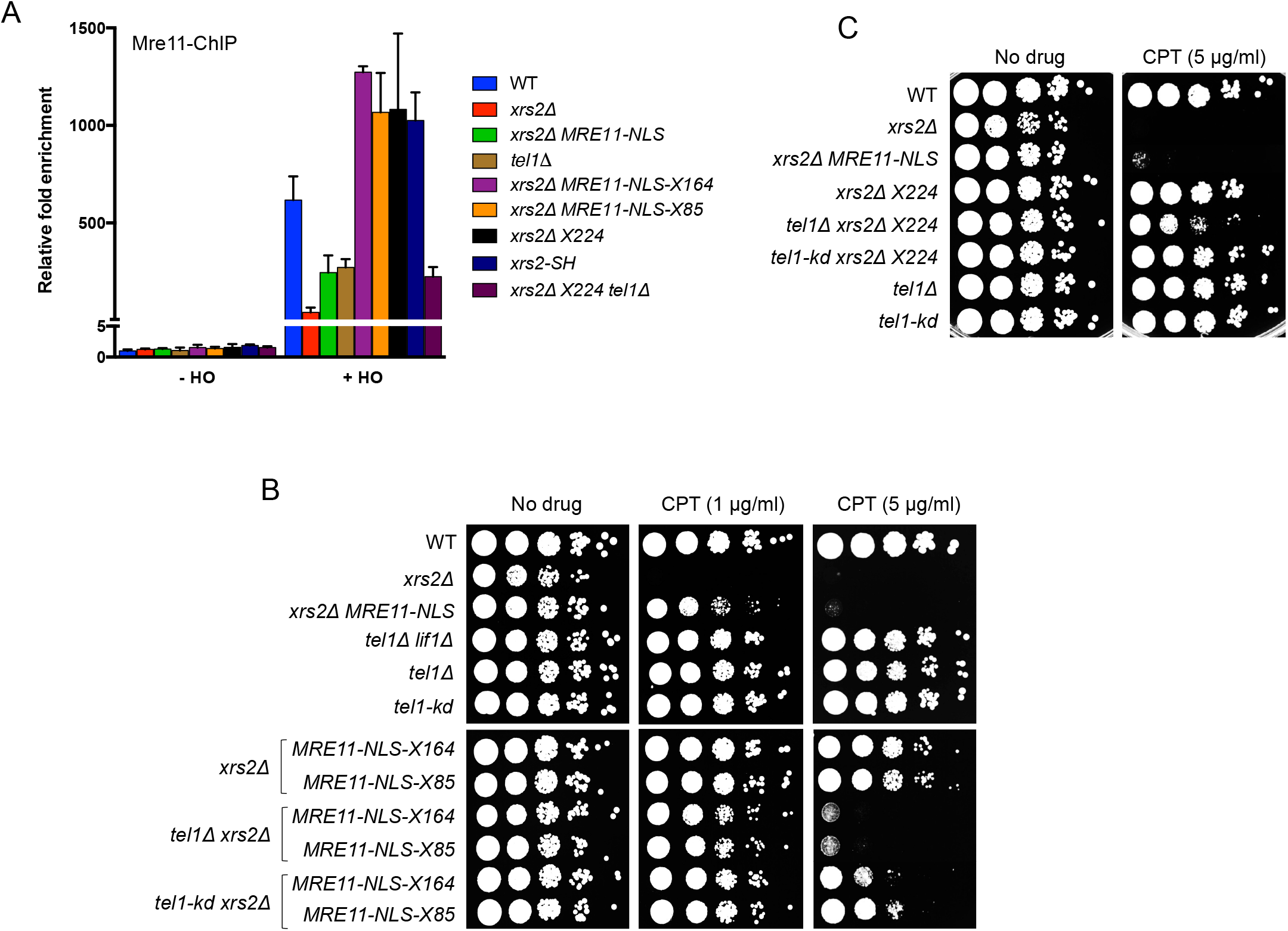
Tel1 promotes stable binding of Mre11 to DSBs and enhances DNA damage resistance. (A) ChIP-qPCR for Mre11 0.2 kb from the HO-induced DSB. The error bars indicate SD (n = 3). (B) and (C) 10-fold serial dilutions of the indicated strains spotted onto rich medium with or without CPT at indicated concentrations.

The severe genotoxin sensitivity of the *xrs2Δ* mutant is partially suppressed by *MRE11-NLS* (Oh et al., 2016; Tsukamoto et al., 2005). However, at a higher concentration of camptothecin (CPT) or methyl methanesulfonate (MMS), the *MRE11-NLS xrs2Δ* strain shows greatly reduced survival as compared to WT (Fig 2B and Fig S2A). This result is not due to the combined defects in Tel1 signaling and NHEJ since the *lif1Δ tel1Δ* double mutant is more resistant to CPT than the *MRE11-NLS xrs2Δ* strain. We hypothesized that the reduced DNA damage resistance of *MRE11-NLS xrs2Δ* cells could be due to failure to maintain Mre11 at DSBs. Indeed, the *MRE11-NLS-TID xrs2Δ* strains, which show restored Mre11 binding to DSBs, exhibit similar DNA damage resistance to WT cells. The restoration of CPT resistance is dependent on Tel1 but is partially independent of the Tel1 kinase activity, consistent with Tel1 contributing in a structural manner to stabilize Mre11 at DSBs. Since deletion of *TEL1* confers CPT sensitivity only in the absence of *XRS2*, it suggests that Tel1 can compensate for Xrs2 in promoting DNA damage resistance (Fig 2B).

The *X224* fragment also restores DNA damage resistance to WT levels; however, unlike the fusion proteins, the restoration is mostly independent of Tel1 (Fig 2C). This result suggests that the Mre11 binding domain present in the 224 aa Xrs2 fragment, but not in the fusion proteins, promotes DNA damage resistance in the absence of Tel1. In agreement with our previous study, normal growth and DNA damage resistance of the *xrs2Δ* strains expressing either the Mre11-TID fusions or X224 fragment is dependent on *SAE2* (Fig S2B,C), indicating that the MR end resection function is critical for proliferation and DNA damage resistance in Xrs2-deficient cells.

### Tel1 rescues the DNA bridging defect of *MRE11-NLS xrs2Δ* cells

The finding that restoring Mre11 enrichment at DSBs enhances DNA damage resistance of the *xrs2Δ* mutant prompted us to examine the structural role of the MRX complex in bridging DSB ends and tethering sister chromatids. In vitro, Mre11-Rad50 is sufficient for end-bridging activity (Deshpande et al., 2014), and the role of Xrs2 in this process has not been investigated. To monitor DSB tethering, we inserted lacO and tetO arrays on opposite sides of an I-SceI cut site on chromosome V of haploid cells (Fig 3A). In this strain, I-SceI is expressed from a galactose-inducible promoter, LacI-YFP and TetR-RFP are constitutively expressed, and a Rad52-CFP fusion is used to monitor DSB formation. It is important to note that the DSB is effectively “irreparable”: HR cannot be employed because I-SceI is expected to cut both sister chromatids in S/G2 phase cells, and imprecise NHEJ to mutate the I-SceI cut site is rare in yeast (Deng et al., 2014).

**Figure 3.**
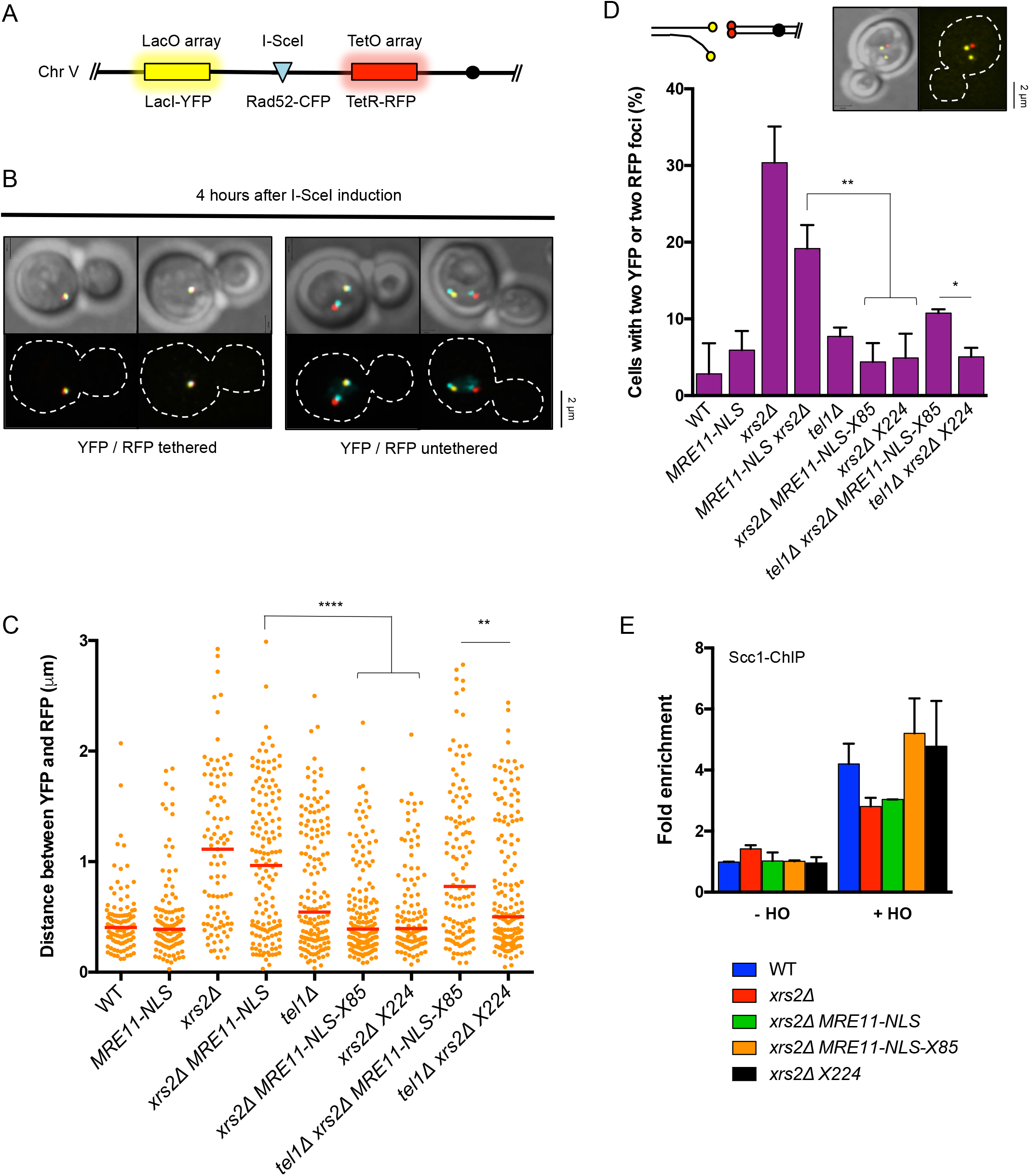
Tel1 promotes the DNA bridging function of the Mre11-Rad50 complex. (A) Schematic representation of the DSB end-tethering assay system. (B) Examples of cells with YFP and RFP foci that are together or separated 4 hr after I-SceI induction. (C) Distribution of the distance between YFP and RFP foci. Red lines indicate median values. Cells in G2/M with Rad52 foci were scored (n ≥ 100). ** P ≤ 0.001, **** P ≤ 0.0001. (D) Cartoon of sister-chromatid separation after DSB formation and image of a cell with two YFP foci, indicating sister-chromatid separation. Graph shows the percentage of cells with either two YFP or two RFP foci. Cells in G2/M were scored (n ≥ 100). * P ≤ 0.05, ** P ≤ 0.01. (E) ChIP-qPCR for Scc1-PK 1 kb from the HO-induced DSB. The error bars indicate SD (n=2).

Four hours after I-SceI induction, the DSB can be visualized by appearance of a Rad52-CFP focus that co-localizes with YFP and/or RFP. For a tethered DSB we cannot distinguish between one and two Rad52 foci (Fig 3B). However, for untethered DSBs we observe some cells with two Rad52 foci, each associated with RFP or YFP, and others with a Rad52 focus associated with only one end (Fig 3B, S3A). The distance between YFP and RFP foci was measured in at least 100 cells with co-localizing Rad52 foci. Consistent with previous studies using similar assays, most WT cells exhibit co-localizing YFP and RFP foci (Fig 3B, C) (Cassani et al., 2016; Kaye et al., 2004; Lobachev et al., 2004; Seeber et al., 2016). We observed a significant increase in DSB end-to-end separation in *xrs2Δ* cells and *MRE11-NLS* is unable to rescue this defect. In agreement with previous studies, the *tel1Δ* mutant shows a slight increase in DSB end separation (Cassani et al., 2016; Lee et al., 2008), but in contrast to the Lee et al study (Lee et al., 2008), we find comparable end tethering in *tel1-kd* and WT cells (Fig S3B). Retention of Mre11 at DSBs is independent of Tel1 kinase activity (Gobbini et al., 2015), correlating with the end-tethering function. As expected if Mre11 retention at DSBs facilitates end tethering, the Mre11-NLS-X85 fusion protein and Xrs2 fragment are able to significantly rescue the end-tethering defect of *xrs2Δ* cells (Fig 3C). The recovery of end tethering in *xrs2Δ MRE11-NLS-X85* cells is Tel1 dependent, while loss of Tel1 in *xrs2Δ X224* cells reduces end tethering to the same level as observed in the *tel1Δ* mutant. These data mirror the DNA damage resistance of the strains and indicate separable roles of Xrs2 binding to Mre11 and Tel1-mediated stabilization of Mre11 DNA association in promoting end tethering and genotoxin resistance. The enhanced end tethering in *xrs2Δ MRE11-NLS-X85* and *xrs2Δ X224* is partially dependent on the kinase activity of Tel1 (Fig S3) suggesting that Tel1 contributes in both a kinase-dependent and -independent manner in these mutant contexts.

In late S and G2 phases, when the sister chromatid is present, the MRX complex also holds sisters together at DSBs (Seeber et al., 2016). Cells with two foci of the same fluorescence indicate sister-chromatid separation. The *xrs2Δ* and *MRE11-NLS xrs2Δ* strains show increased sister chromatid separation, which again is rescued by the fusion proteins and *X224* fragment (Fig 3D). These data demonstrate that Tel1 recruitment is crucial to stabilize Mre11 at DSBs to facilitate the DNA bridging function of the complex, especially when Xrs2 is not present. Cohesin, an SMC complex that normally keeps sister chromatids paired during G2 and cell division, also contributes to DSB and stalled replication fork repair, presumably by maintaining sister chromatids in a conformation that favors HR (Heidinger-Pauli et al., 2008; Kim et al., 2002; Sjogren and Nasmyth, 2001; Tittel-Elmer et al., 2012). Mre11 and Tel1 are involved in recruitment of DNA damage-induced cohesin around DSBs and stalled forks (Strom et al., 2007; Strom and Sjogren, 2007; Tittel-Elmer et al., 2012; Unal et al., 2004; Unal et al., 2007). To assess cohesin binding, enrichment of Scc1 (one of the subunits of the cohesin complex) was measured at sequences 1 kb from an HO induced DSB at the *MAT* locus. The reduced Scc1 binding observed in *xrs2Δ* and *xrs2Δ MRE11-NLS* cells is rescued by expression of the fusion protein as well as *X224* fragment (Fig 3E). This observation suggests that the sister chromatid separation in *xrs2Δ* cells could be due to reduced cohesin recruitment resulting from low enrichment of Mre11 at DSB ends.

### End tethering by MRX is not required for DSB-induced recombination

Previous studies have suggested that the end tethering function of MRX is important for NHEJ and HR (Cassani et al., 2018; Cassani et al., 2016; Chen et al., 2001; Deshpande et al., 2014). We tested whether NHEJ is restored in *xrs2Δ* cells expressing the fusion proteins or *X224* fragment since end-to-end tethering is significantly increased in these cells. Using a plasmid-ligation assay we found that NHEJ is at the same low level in all of the *xrs2Δ* derivatives (Fig S4A), indicating that restoration of end tethering is not sufficient for NHEJ. Previous studies have shown that interaction between the Xrs2 FHA domain and Lif1 is required for NHEJ, and is likely the reason for low NHEJ in the *xrs2Δ* strains expressing *MRE11-NLS-TID* or *X224* (Chen et al., 2001; Matsuzaki et al., 2008; Oh et al., 2016; Palmbos et al., 2008).

Next, we used a direct repeat recombination reporter to determine how end tethering affects DSB-induced HR. In this system, an I-SceI induced DSB at the *ade2-I* locus is repaired using the intact *ade2-n* allele (Fig 4A) (Mozlin et al., 2008). In *RAD51* cells, repair occurs mainly by gene conversion (GC) maintaining the *TRP1* marker located between the repeats, whereas single strand annealing (SSA), which results in deletion of *TRP1* and one of the repeats, is *RAD51*-independent. We observe no significant change in *RAD51*-dependent GC or *RAD51*-independent SSA in *MRE11-NLS xrs2Δ* compared to WT (Fig. 4B), indicating that end tethering is not required for homology-dependent DSB repair in this context.

**Figure 4.**
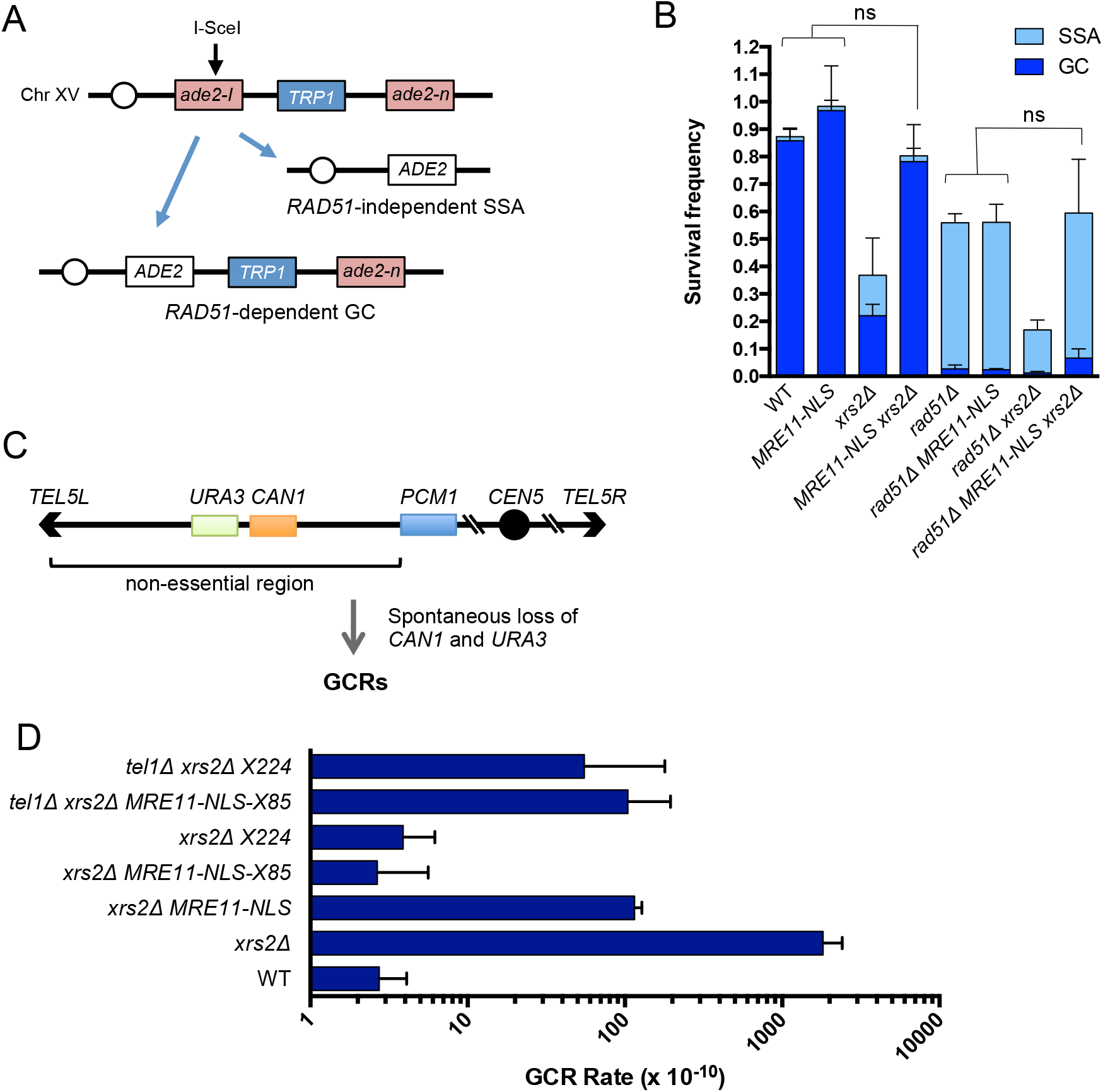
End tethering is required to suppress GCRs. (A) Schematic of the *ade2* direct repeat recombination reporter. Repair of the I-SceI induced DSB occurs mostly by gene conversion with no accompanying crossover retaining *TRP1*. SSA, a *RAD51*-independent process, results in loss of one of the repeats and the intervening *TRP1* marker. (B) The frequencies of DSB-induced GC and SSA repair for the indicated strains. Error bars indicate SD (n=3). ns = not significant (P ≥ 0.05). (C) Schematic of the GCR assay. Simultaneous loss of *URA3* and *CAN1* (selected by growth of cells on medium containing 5-FOA and canavanine) is due to loss of the terminal chromosome region followed by telomere addition, interstitial deletion, non-reciprocal translocation of hairpin-mediated inverted duplication. (D) GCR rates measured by simultaneous loss of *CAN1* and *URA3*. Error bars indicate 95% confidence interval (n ≥ 10).

Null mutation of genes encoding the MRX complex results in an increased rate of spontaneous recombination between heteroalleles in diploid cells (Ajimura et al., 1993; Ivanov et al., 1992; Malone et al., 1990). One mechanism suggested for the hyper-recombination phenotype is by channeling lesions from the sister chromatid to the homologue for repair due to disruption of sister-chromatid tethering (Hohl et al., 2015; Symington et al., 2014). We measured the rate of spontaneous recombination using diploid cells with *ade2-I* and *ade2-n* heteroalleles (Fig S4B). Consistent with a previous study, the *xrs2Δ* mutant displays a 5-fold increase in the rate of Ade^+^ recombinants (Ivanov et al., 1992). Surprisingly, this phenotype is suppressed by *MRE11-NLS*, indicating that defective sister chromatid tethering is not responsible for the hyper-recombination phenotype (Fig S4C). However, when *SAE2* is deleted in this strain, the triple mutant again shows hyper-recombination suggesting that the hyper-rec phenotype is due to defective end resection, reducing co-conversion of the markers.

### Suppression of chromosome rearrangements by MRX

Previous studies reported a 600-fold increase in the rate of gross chromosome rearrangements (GCRs) in the absence of MRX (Chen and Kolodner, 1999). By contrast, loss of Tel1 signaling or Mre11 nuclease activity causes no increase or modest increase in GCRs, respectively (Deng et al., 2015; Myung et al., 2001; Smith et al., 2005). We measured the spontaneous GCR rate using an assay that detects simultaneous loss of two markers on the left arm of chromosome V (Chen and Kolodner, 1999) (Fig 4C). Consistent with previous studies, the *xrs2Δ* mutant shows a 664-fold increase in GCR accumulation compared to WT. *MRE11-NLS* lowers the GCR rate of *xrs2Δ* cells ~16-fold, but this rate is still ~42-fold higher than observed for WT (Fig 4D). However, expression of the *MRE11-NLS-X85* fusion protein or *X224* fragment in the *xrs2Δ* mutant led to a complete suppression of the hyper-GCR phenotype in a Tel1 dependent manner.

### Tel1 rescues the stalled replication fork instability of *MRE11-NLS xrs2Δ* cells

We noticed a significant increase in cells with spontaneous Rad52-CFP foci in *xrs2Δ* and *MRE11-NLS xrs2Δ* strains (Fig 5A). This observation, along with the increased rate of GCRs and sensitivity to hydroxyurea (HU) (Fig 5B), suggests more replication-associated DNA damage. The MRX complex is recruited to stalled replication forks and has been shown to stabilize the association of essential replisome components (Seeber et al., 2016; Tittel-Elmer et al., 2009). This function is independent of the S-phase checkpoint and the nuclease activity of Mre11, indicating a structural contribution of the complex in stabilizing stalled replication forks (Tittel-Elmer et al., 2009). To address whether the DNA bridging function of MRX correlates with the replisome stability function, we measured the presence of Mre11 and DNA polymerase α (Polα) near an early firing origin (*ARS607*) by ChIP after releasing G1 synchronized cells into 0.2 M HU. As anticipated, the strains with DNA tethering defects, *xrs2Δ* and *MRE11-NLS xrs2Δ*, show loss of Mre11 and Polα enrichment compared to WT (Fig 5C). Consistently, expression of *MRE11-NLS-X85*, as well as the *X224* fragment, completely rescues Mre11 and Polα enrichment at stalled replication forks and HU sensitivity of *xrs2Δ* cells, as well as reducing the number of cells with spontaneous Rad52 foci (Fig 5A, B, C). Unlike the response to CPT and MMS, we find HU resistance of the *xrs2Δ X224* mutant requires Tel1. Therefore, we also measured Polα and Mre11 enrichment at *ARS607* in *tel1Δ* derivatives. Polα enrichment in *tel1Δ* cells is comparable to WT cells, while Mre11 enrichment is reduced, similar to that observed at DSBs. The rescue of Polα enrichment is completely Tel1-dependent in *xrs2Δ MRE11-NLS-X85* cells (Fig 5C), consistent with the end tethering data, suppression of spontaneous Rad52 foci and GCRs. At the 40 min time point, Polα retention at *ARS607* in *xrs2Δ X224* cells is partially Tel1 dependent, but at 60 min, Polα enrichment is lost in the *tel1Δ* derivative, correlating with GCR results. These data suggest that Tel1 stabilization of Mre11 at stalled forks is important to prevent fork collapse and suppression of GCRs in cells lacking Xrs2.

**Figure 5:**
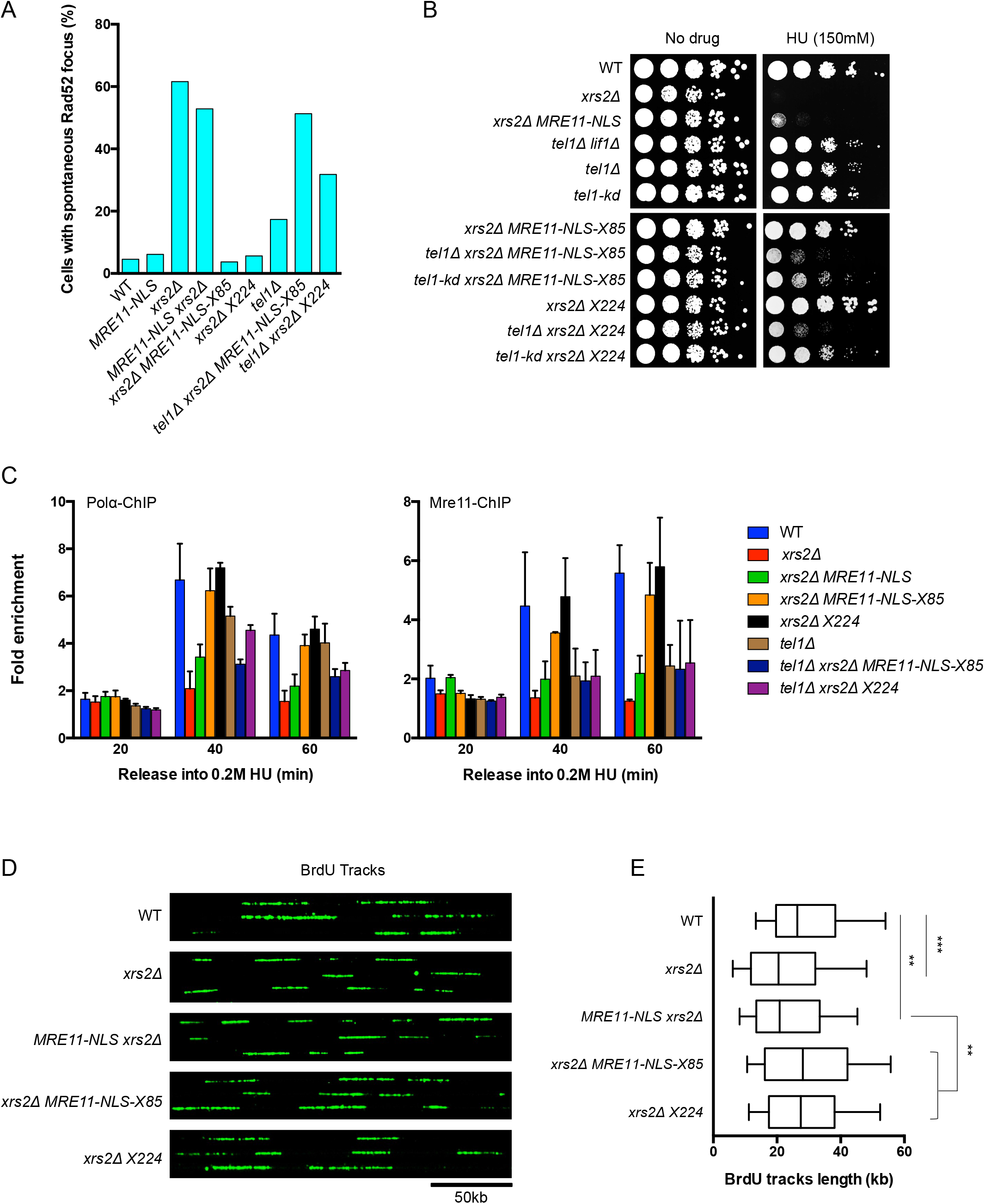
Tel1 promotes the stalled-replication fork stability in *MRE11-NLS xrs2Δ* cells. (A) Percentage of cells with a spontaneous Rad52 focus. (n ≥ 100). (B) 10-fold serial dilutions of the indicated strains spotted onto YPD medium with or without HU at indicated the concentration. (C) ChIP-qPCR for Polα-Flag and Mre11 at early firing origin *ARS607* following release of cells from G1 arrest into medium containing 0.2 M HU. The error bars indicate SD (n ≥ 2). (D) Representative DNA fibers after combing and detection of BrdU tracks in green. Genomic DNA was obtained from S phase cells labeled for 3 hr with BrdU in the presence of 0.2 M HU. (E) Distribution of BrdU track lengths in HU-treated cells. Box: 25-75-percentile range. Whiskers: 10-90-percentile range. Vertical bars indicate median values. ** P ≤ 0.01, *** P ≤ 0.001 (Mann-Whitney rank sum test).

To visualize replication fork progression in the presence of replicative stress, DNA combing was performed. Genomic DNA obtained from S phase cells pulse labeled with BrdU for 3 hr in the presence of 0.2 M HU were stretched and newly synthesized DNA tracks were detected with anti-BrdU (Fig 5D). Consistent with the DNA Polα ChIP data, *xrs2Δ* and *MRE11-NLS xrs2Δ* had shorter track lengths compared to WT, which was rescued by the *MRE11-NLSX85* fusion protein or the *X224* fragment (Fig 5E). Collectively, these data show that loss of MR-mediated end tethering correlates with increased replisome fragility and elevated genomic instability.

## Discussion

MRX functions in telomere maintenance and DNA damage checkpoint signaling by recruiting and activating the Tel1 kinase. Although the Tel1 binding domain within the C-terminal region of yeast Xrs2/Nbs1 is required for Tel1 activation in vivo (Nakada et al., 2003; You et al., 2005), ATM activation can occur independently of the C-terminal ATM-interaction domain of mammalian Nbs1 (Kim et al., 2017; Lee and Paull, 2005; Stracker et al., 2007). Moreover, ATP-induced conformational changes to the MR complex are critical for ATM activation (Al-Ahmadie et al., 2014; Deshpande et al., 2014; Morales et al., 2005; Roset et al., 2014). Here, we show that fusing the Tel1 interaction domain from Xrs2 to Mre11-NLS restores Tel1 activation, supporting the hypothesis that Tel1 recruitment and activation are separate functions of the MRX complex. Our studies further refine the Tel1 binding domain to the last 84 amino acids of Xrs2, encompassing an acidic patch and FXF/Y motif that were previously shown to be essential for Tel1/ATM signaling in other systems (Falck et al., 2005; Limbo et al., 2018; You et al., 2005).

Unexpectedly, our studies identified a structural role for Tel1 in maintaining Mre11 association with DSBs that becomes physiologically relevant in the absence of Xrs2. Previous studies have shown reduced association of Mre11 with DSBs in the *tel1Δ* mutant, but not in cells lacking Tel1 kinase activity (Cassani et al., 2016; Gobbini et al., 2015; Oh et al., 2016). The *tel1Δ* mutant shows far greater resistance to genotoxins than *mre11Δ*, indicating that reduced binding of Mre11 at damage sites does not grossly impair the DNA repair function of Mre11. Nbs1 binding to Mre11 extends the dimer interface and stabilizes the dimeric form of Mre11 (Schiller et al., 2012). We find that expression of an Xrs2-derived peptide encompassing the Mre11 and Tel1 binding domains is highly effective in suppressing DNA damage resistance and Mre11 retention at DSBs in *xrs2Δ* cells, suggesting that stabilization of the Mre11 dimer is a critical function of Xrs2. Supporting our findings, Kim et al (2017) found that expression of a 108 amino acid fragment of Nbs1, encompassing the Mre11 interaction domain, is sufficient to restore proliferation to Nbs1-deficient murine cells. Our data suggest that Tel1 and Xrs2 independently contribute to Mre11 activity at DNA ends. While loss of Tel1 stabilization alone does not have a strong impact on DNA damage resistance, Tel1 can compensate for Xrs2-mediated Mre11 dimer stabilization to promote repair. The Tel1 stabilization function is critical for HU resistance and suppression of GCRs, even when the Xrs2-Mre11 interaction interface is restored, suggesting an additional function of Xrs2 during replication stress. Interestingly, the murine *Nbs1^ΔB/ΔB^* mutation, which deletes the N-terminal FHA and BRCT domains but retains Mre11 interaction, is synthetically lethal with ATM deficiency, suggesting that compensation between Nbs1 and ATM is conserved in mammals (Williams et al., 2002). We propose that the quantity and quality of the MRX complex compensate each other. Optimally stabilized Mre11 complex may engage in sufficient DNA tethering with minimal quantity while suboptimal complex may exhibit reduced ability interact with DNA and thus require a higher local concentration. It remains unclear how Tel1 facilitates Mre11 retention at DSBs because no direct interaction between MR and Tel1 has been reported.

Our findings indicate that Mre11 stabilization at ends is critical for the end tethering function of MRX, and the previously reported reduction in end tethering of the *tel1Δ* mutant is a consequence of lower retention of Mre11 at DSBs. Retention of Mre11 at DSBs, end tethering and DNA damage resistance are highly correlated, raising the question of how end tethering facilitates genome integrity. Although end tethering is restored in *xrs2Δ MRE11-NLS-X85* cells, NHEJ remains defective indicating that end tethering is not sufficient for this mode of repair. Previous studies have suggested that end tethering is important for DSB-induced gene conversion and for SSA (Cassani et al., 2018; Cassani et al., 2016; Ferrari et al., 2015). However, we found both gene conversion and SSA to be restored to wild-type frequencies in *xrs2Δ MRE11-NLS* cells, which are defective for end tethering. Because the assay we used measures intra-chromatid recombination or recombination between misaligned sister chromatids, we cannot rule out the possibility that MRX bridging is important for precise sister chromatid recombination.

Our data suggest that CPT and MMS sensitivity of *xrs2Δ MRE11-NLS* cells is due to failure to maintain tethering during DNA replication. Loss of DNA tethering results in more spontaneous Rad52 foci and increased rates of GCRs, indicators of replication stress. The strains with reduced end tethering show lower Mre11 association with stalled replication forks,replisome instability and shorter DNA synthesis tracks in response to replication stress. This phenotype could be caused by loss of cohesin since a previous study showed that *rad50* mutants defective for tethering have reduced cohesin bound at stalled replication forks (Tittel-Elmer et al., 2012). Consistently, we show that cohesin enrichment mirrors Mre11 enrichment at DSBs. The MR complex is known to associate with chromatin during S-phase and co-localizes to stressed and unstressed replication forks (Mirzoeva and Petrini, 2003; Sirbu et al., 2011; Tittel-Elmer et al., 2009). MR could use its intrinsic DNA binding activity to travel with the replisome, associate with the end produced by fork reversal or could be indirectly associated with DNA via RPA interaction (Seeber et al., 2016).

## Materials and methods

### Media and Growth Conditions

Media and growth conditions were as described previously (Amberg, 2005). Experiments were carried out with log-phase cells, unless otherwise indicated. Cells were grown at 30°C for all the experiments except for the end tethering assay, in which cells were grown at 23°C.

### Yeast strains and plasmids

Yeast strains used in this study are listed in Table S1. For the Mre11-NLS-X164 and Mre11-NLS-X85 strains, overlapping PCR was performed to fuse C-terminal 164 amino acid and 85 amino acid fragments of Xrs2 to the C-terminus of Mre11-NLS with 5xGly as a linker. The resulting Mre11-NLS-X164 and Mre11-NLS-X85 constructs were cloned into pRG205MX (Gnugge et al., 2016), along with the promoter (450bp upstream) and 3’UTR (599bp downstream) of *MRE11*, and were integrated into the *LEU2* locus of *mre11::TRP1* strain. Integration was confirmed by PCR and DNA sequence analysis, and expression of the fusion proteins was analyzed by western blot using α-Mre11 polyclonal antibodies (Krogh et al., 2005). For the X224 strain, a 224 amino acid C-terminal fragment of Xrs2 was amplified and cloned into pRG205MX, along with the *XRS2* promoter (504bp upstream) and 3’UTR (468bp downstream), and was integrated into the LEU2 locus of an *xrs2::kanMX* strain. X224-MYC strain was constructed by one-step targeting of a PCR fragment containing a sequence encoding 13 repeats of MYC. For the end tethering assay strain (W11278-23B), a tandem array of Tet operators (TetOx336) was integrated in *ura3* on chromosome V, and an I-*Sce*I cut site was integrated 4 kb to the left of *ura3* in iYEL023, as described (Lisby et al., 2003). The TetO array was visualized by TetR-mRFP, which is integrated in the intergenic region iYGL119 on chromosome VII. A tandem array of Lac operators (LacOx256) was integrated in iYEL024, 4 kb to the left of the I-*Sce*I cut site. The LacO array was visualized by YFP-LacI integrated in *his3*. The I-*Sce*I endonuclease gene is under the transcriptional control of *GAL1-10* promoter, integrated in *lys2*. The Rad52 C-terminus was tagged with GAx3 linker and yeast codon optimized mTurquoise2, which is an improved variant of CFP. The protein sequence of mTurquoise2 is from (Goedhart et al., 2012) and yeast codon optimization was ordered from GenScript. Integration was accomplished with pop-in, pop-out using *K. lactis URA3*. All other W303-derived strains were obtained from crossing appropriate haploid strains. Strains used for the GCR assay were made by transformation with linear DNA fragments to generate gene disruptions, or to integrate Mre11 or Xrs2 constructs.

### DNA damage sensitivity assays, ChIP-qPCR, western blot, telomere blot, and Rad53 phosphorylation assay

These assays were performed as described (Oh et al., 2016). For Polα-Flag and Mre11 ChIP at replication origin ARS607, cells were arrested in G1 using α-factor and released into YPD containing 0.2M HU. Cells were collected at 20 min intervals after release. Anti-Mre11 (Krogh et al., 2005), anti-HA (ab9110), anti-Flag M2 (Sigma), and anti-V5 [SV5-pk1] (ab27671) were used for ChIP-qPCR. qPCRs were carried out by the SYBR green system using primer pairs complementary to DNA sequences corresponding to *ARS607* and sites 0.2 kb and 1 kb from the HO-cut site at *MAT*. DNA sequences located 66 kb from *MAT* and 14 kb from ARS607 were amplified as controls. Fold enrichment calculations were done as described (Oh et al., 2016).anti-Mre11, anti-Rad53 (gift from M. Foiani), and anti-cMyc 9E10 (Santa Cruz Biotechnology) were used for western blots.

### End tethering assay

Cells were grown to log phase in SC medium containing 2% raffinose at 23°C. Then, galactose was added to a final concentration of 2% for I-SceI induction. Cells were collected and washed after 4 hr of growth at 23°C. Cells were re-suspended in a small volume of SC medium containing 2% glucose and were immobilized on a microscope slide by mixing them with a solution of 1.2% low melting agarose in SC medium. Live cell fluorescent imaging was performed on Leica DM5500B upright microscope with 100x Leica oil-immersion 1.46NA objective, using 100W mercury arc lamp as the light source. Chroma bandpass filter sets were used to visualize RFP (41002c), YFP (41028), and CFP (31044v2). Images were acquired with Hamamatsu ORCA-ER-1394 camera using Volocity software. 14 z-sections at 0.3µm intervals were taken for each channel.

### DNA combing

DNA combing was performed as described (Hélène Tourrière, 2017). BrdU was detected with a rat monoclonal antibody (ab6326) followed by a secondary antibody coupled to Alexa 488 (A11006, Molecular Probes). DNA molecules were detected with an anti-ssDNA antibody (MAB3034) followed by an anti-mouse IgG coupled to Alexa 546 (A11030, Molecular Probes). DNA fibers were analyzed on a Zeiss Axio Imager 2 microscope equipped with AxioCam MRc and a 63x Zeiss oil-immersion objective. Image acquisition was performed with AxioVision software. The BrdU track length was measured with ImageJ and representative DNA fibers were assembled with Image J.

### GCR assay

The rate of GCRs was measured by fluctuation assay as previously described (Putnam and Kolodner, 2010). Two or more independent experiments using sets of at least five independent cultures were performed.

### Recombination assays

The direct repeat recombination assay was performed as previously described (Ruff et al., 2016). For the diploid recombination assay, diploids were grown to log phase and plated on YPD or SC-ADE. Plates were incubated at 30°C and counted after 3-4 days. Fluctuation assay was used to determine the heteroallelic recombination rate. Three independent experiments using sets of eight independent cultures were performed.

## Acknowledgements

We thank M. Foiani for Rad53 antibodies, J. Cobb and M. P. Longhese for yeast strains, and W.K. Holloman for critical discussion of the data. This study was supported by grants from the National Institutes of Health (R35 GM126997 and P01CA174653 to L.S.S and R35 GM118180 to R.R.)

## Author contributions

J.O. performed the experiments shown in Figs 1, 2, 4, 5, S1, S2, S4 and contributed to Figs 3 and S3. S.J.L. and R.R. constructed the strain used for the end-tethering assay and contributed to data collection and analysis for Figs 3, Fig 5 and S3. J.O and L.S.S. designed the study and wrote the manuscript.

---

**Figure S1.**
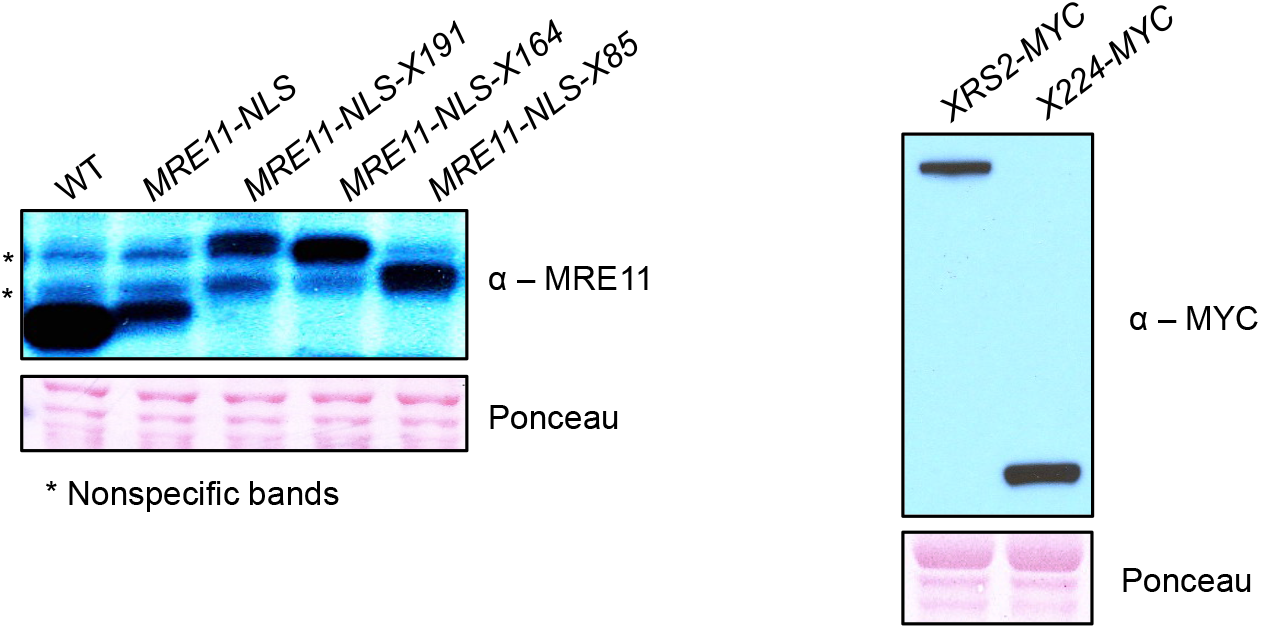
Expression of fusion proteins and X224 peptide, related to Figure 1. Steady-state protein levels of Mre11 and Xrs2 of indicated strains measured by western blot analysis.

**Figure S2.**
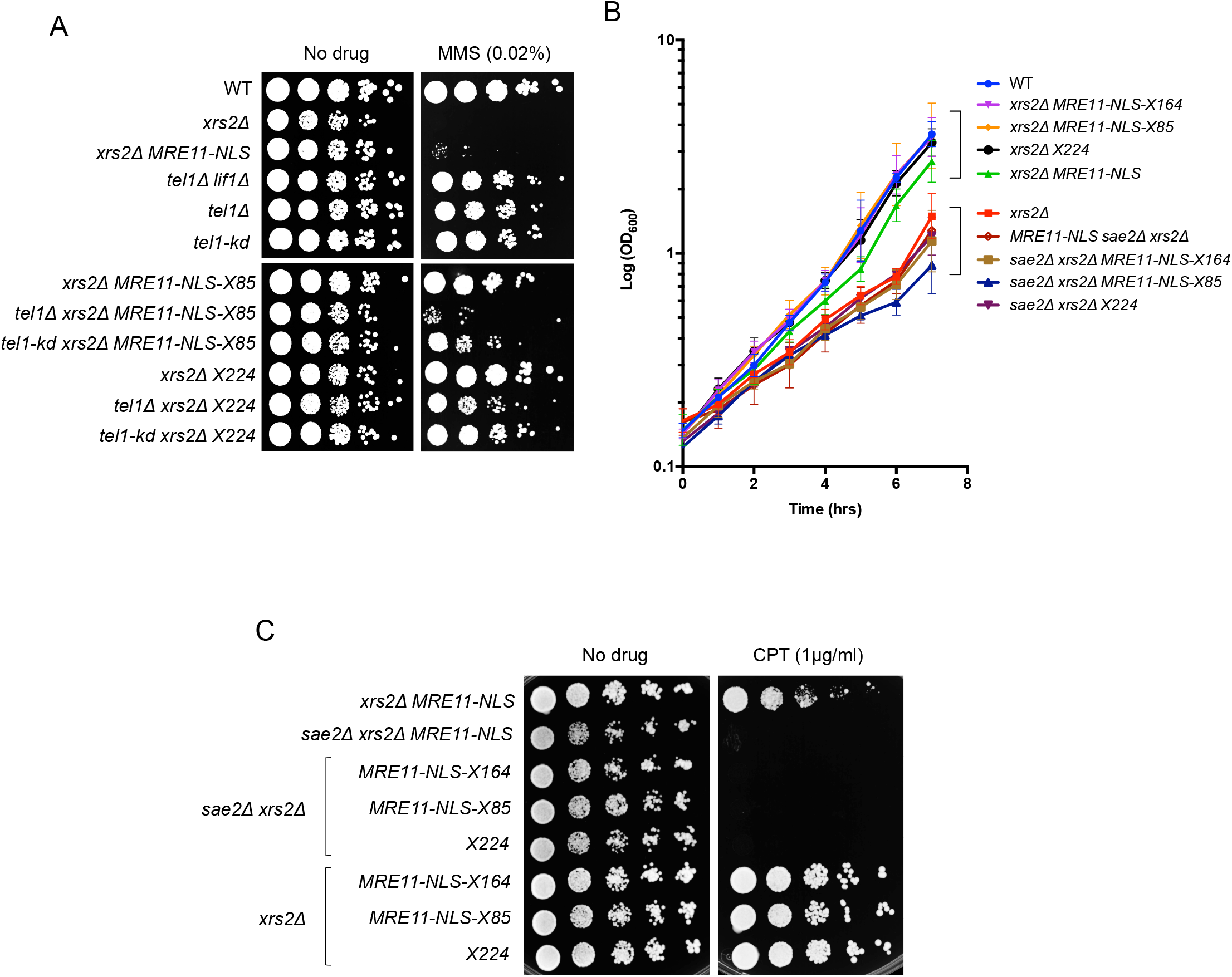
Mre11 fusions and X224 fragment alleviate the MMS sensitivity of *xrs2Δ MRE11-NLS*. Complementation of the growth and DNA damage resistance defects of *xrs2Δ* by the Mre11 fusions and X224 fragment is dependent on *SAE2*, related to Figure 2. A. 10-fold serial dilutions of the indicated strains spotted onto YPD medium with or without MMS at indicated the concentration. B. Growth curves representing cell concentration measured by OD_600_ at the indicated time points. Error bars indicate SD (n=3). C. 10-fold serial dilutions of the indicated strains spotted onto rich medium with our without CPT.

**Figure S3. Related to Figure 3.**
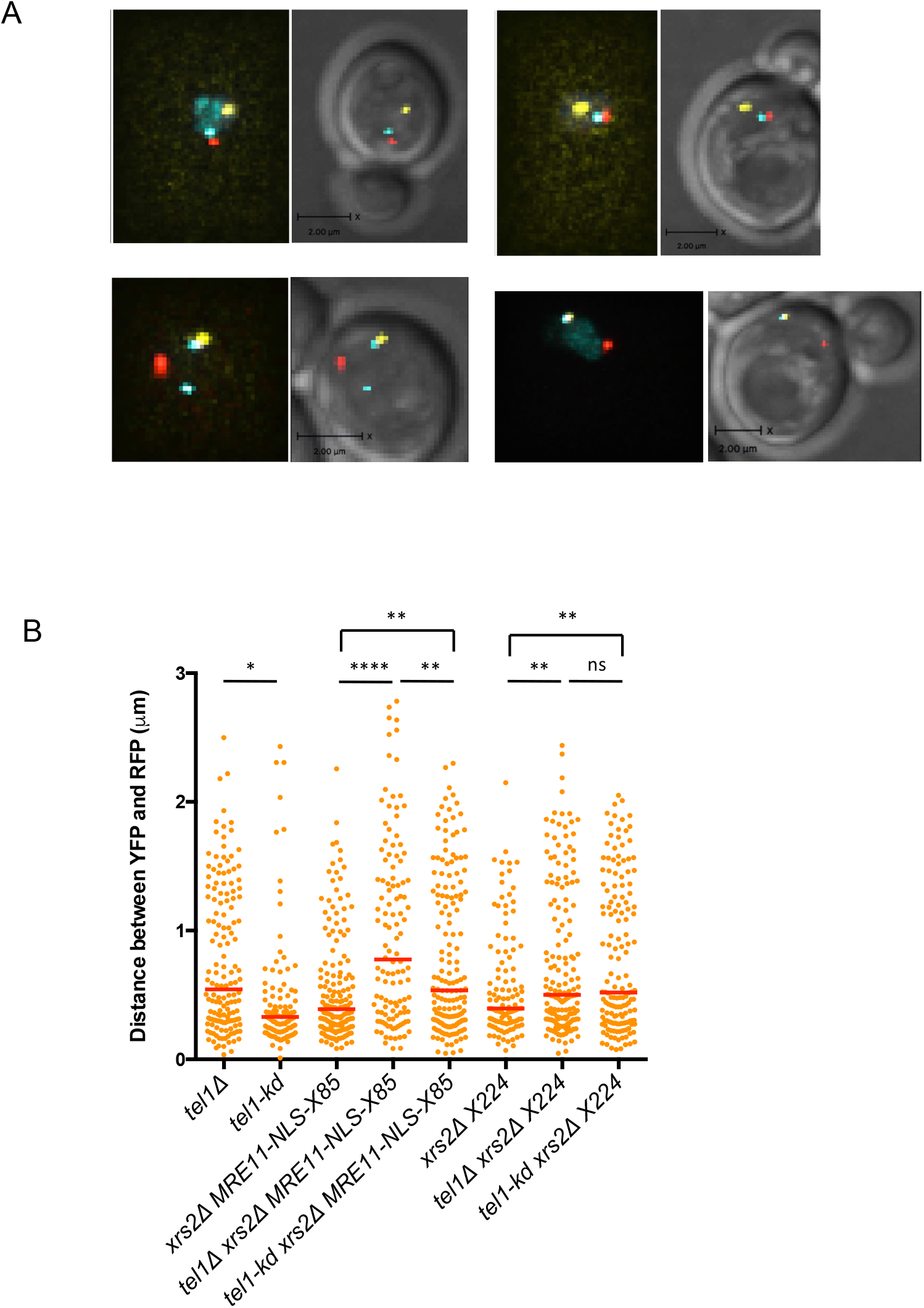
A. Examples of cells with YFP and RFP foci that are separated 4 hr after I-SceI induction. Note that Rad52-CFP is sometimes visible at only one end, and spontaneous Rad52 foci are observed that fail to colocalize with YFP or RFP. B. Distribution of the distance between YFP and RFP foci in *tel1Δ* and *tel1-kd* derivatives.). * P ≤ 0.05, ** P ≤ 0.001, **** P ≤ 0.0001, ns = not significant (P ≥ 0.05).

**Figure S4.**
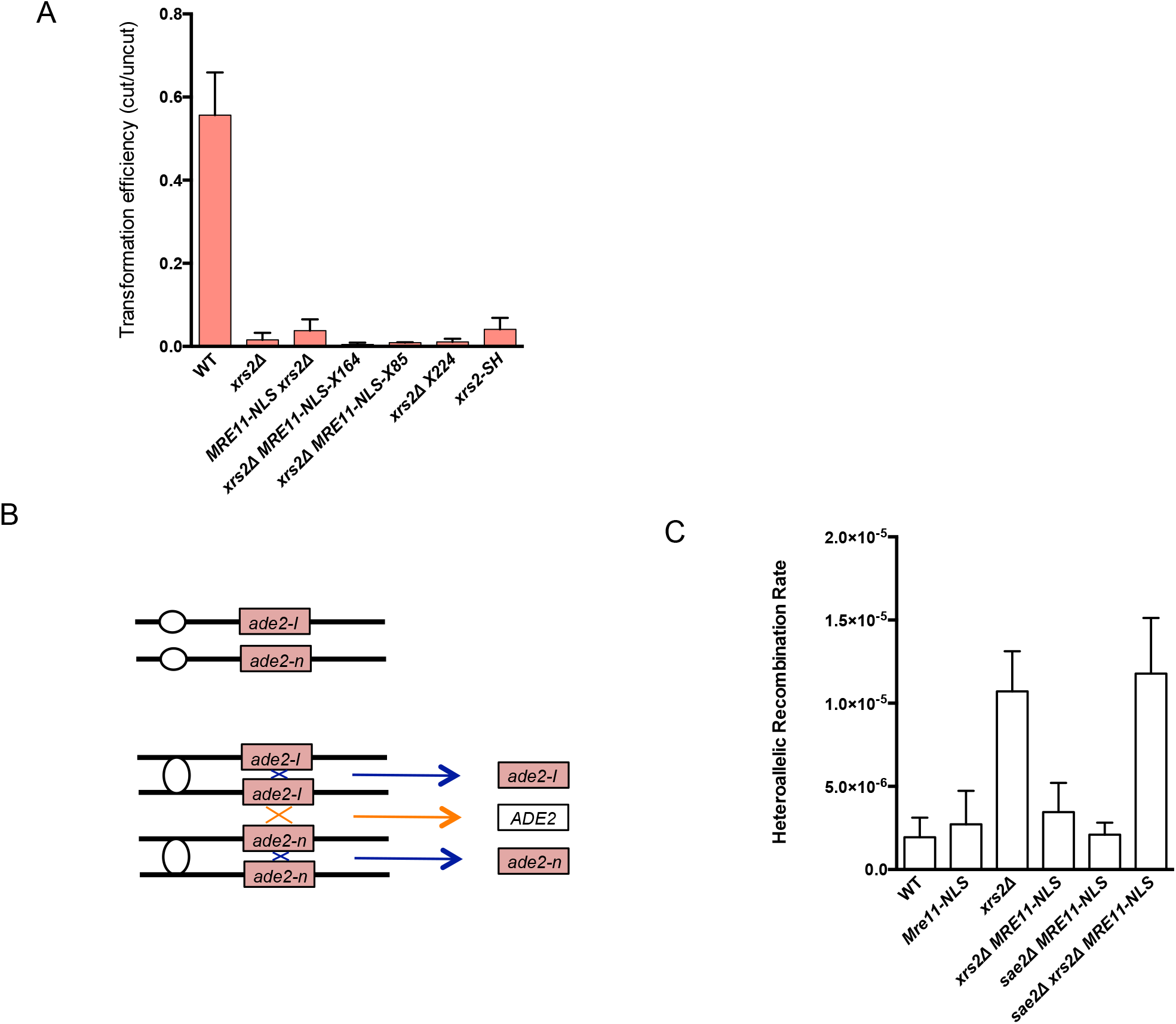
End tethering is not required for NHEJ or HR, related to Figure 4. A. Frequency of NHEJ measured by the ratio of transformants recovered from cut plasmid relative to uncut plasmid for the indicated strains. Error bars indicate SD (n=3). B. Schematic of the diploid recombination assay. In G2 phase cells, spontaneous lesions can be repaired from the sister chromatid resulting in no genetic alteration, or between non-sisters resulting in restoration of *ADE2*. C. Rate of Ade*+* recombinants in the indicated strains. Error bars indicate SD (n=3).

**Supplementary Table 1.**
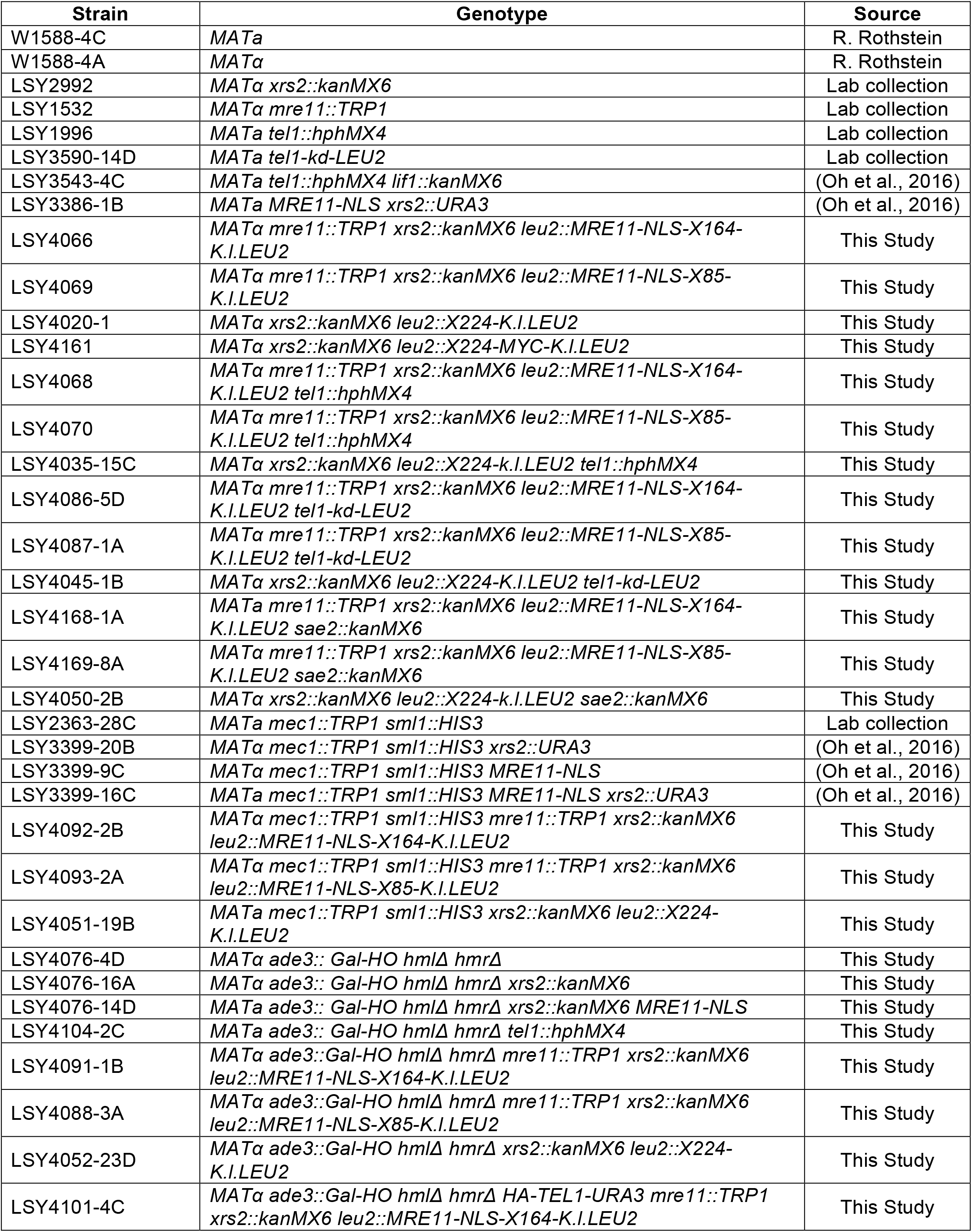
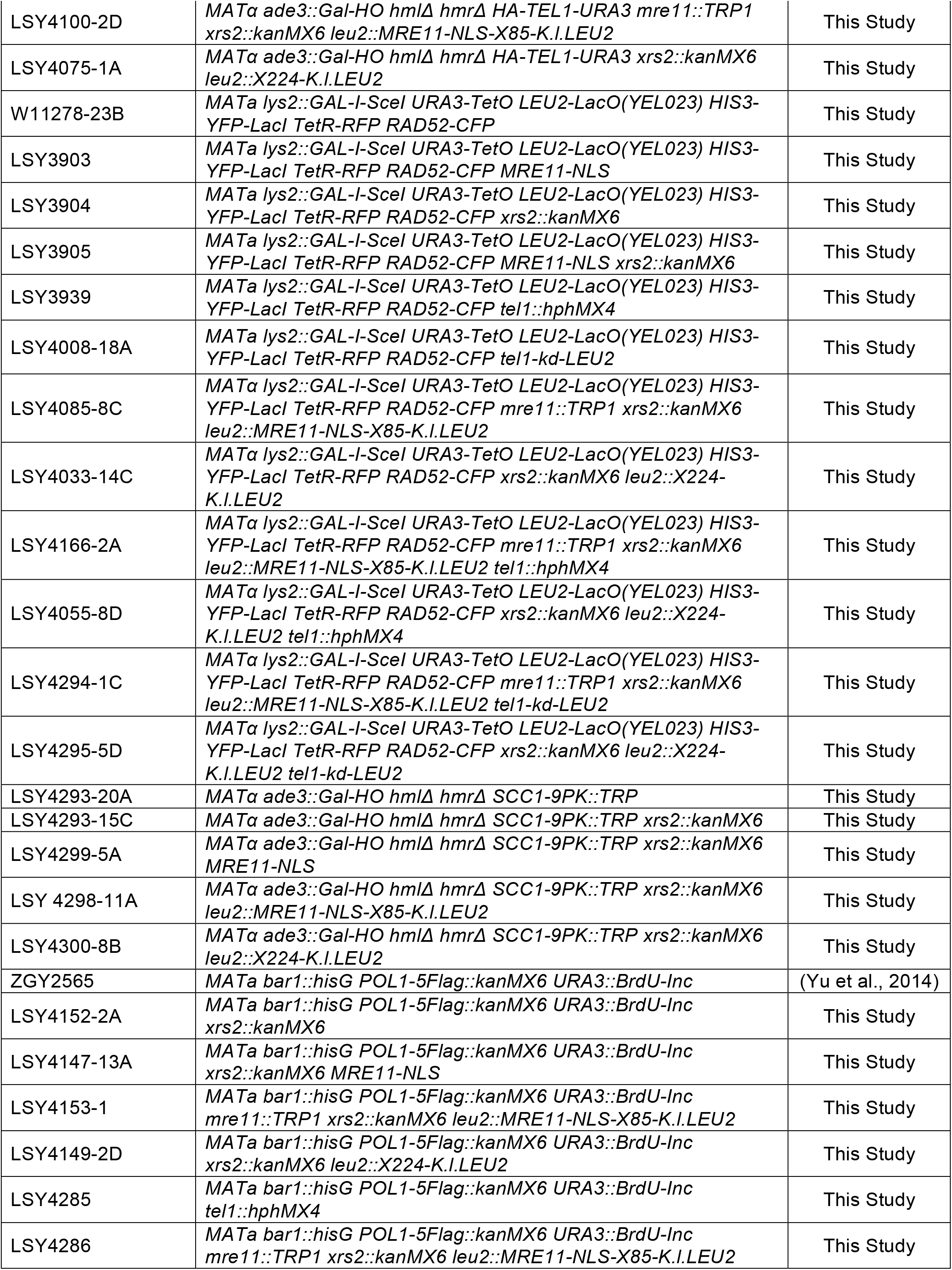
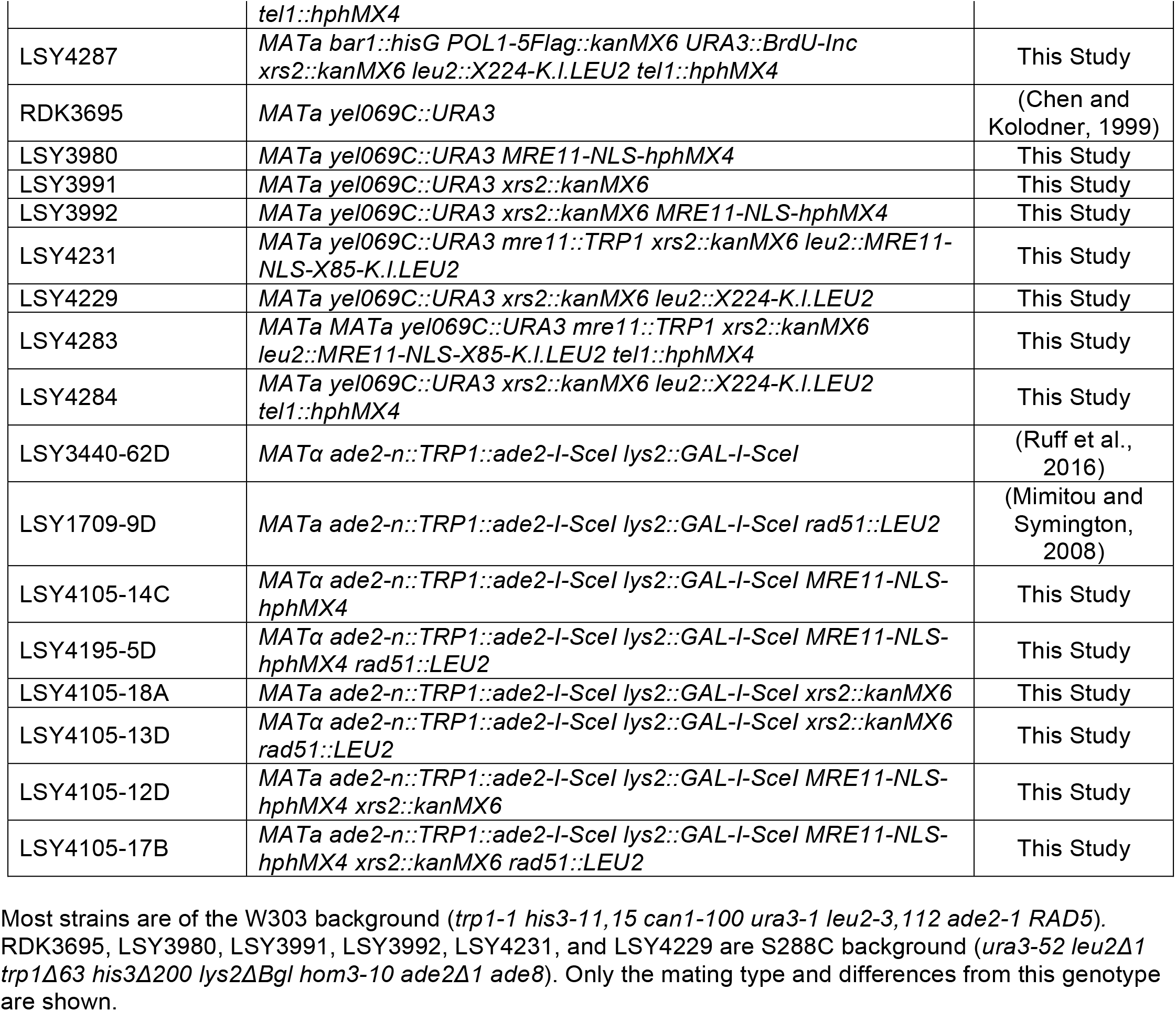

## References

Ajimura, M., Leem, S.H., and Ogawa, H. (1993). Identification of new genes required for meiotic recombination in Saccharomyces cerevisiae. Genetics 133, 51–66.

Al-Ahmadie, H., Iyer, G., Hohl, M., Asthana, S., Inagaki, A., Schultz, N., Hanrahan, A.J., Scott, S.N., Brannon, A.R., McDermott, G.C., et al. (2014). Synthetic lethality in ATM-deficient RAD50-mutant tumors underlies outlier response to cancer therapy. Cancer Discov 4, 1014–1021.

Amberg, D.C., Burke, D. J. & Strathern, J. N. (2005). Methods in Yeast Genetics: A Cold Spring Harbor Laboratory Course Manual. Cold spring Harbor Laboratory Press.

Borde, V. (2007). The multiple roles of the Mre11 complex for meiotic recombination. Chromosome Res 15, 551–563.

Carney, J.P., Maser, R.S., Olivares, H., Davis, E.M., Le Beau, M., Yates, J.R., 3rd, Hays, L., Morgan, W.F., and Petrini, J.H. (1998). The hMre11/hRad50 protein complex and Nijmegen breakage syndrome: linkage of double-strand break repair to the cellular DNA damage response. Cell 93, 477–486.

Cassani, C., Gobbini, E., Vertemara, J., Wang, W., Marsella, A., Sung, P., Tisi, R., Zampella, G., and Longhese, M.P. (2018). Structurally distinct Mre11 domains mediate MRX functions in resection, end-tethering and DNA damage resistance. Nucleic acids research.

Cassani, C., Gobbini, E., Wang, W., Niu, H., Clerici, M., Sung, P., and Longhese, M.P. (2016). Tel1 and Rif2 Regulate MRX Functions in End-Tethering and Repair of DNA Double-Strand Breaks. PLoS Biol 14, e1002387.

Chen, C., and Kolodner, R.D. (1999). Gross chromosomal rearrangements in Saccharomyces cerevisiae replication and recombination defective mutants. Nat Genet 23, 81–85.

Chen, L., Trujillo, K., Ramos, W., Sung, P., and Tomkinson, A.E. (2001). Promotion of Dnl4-catalyzed DNA end-joining by the Rad50/Mre11/Xrs2 and Hdf1/Hdf2 complexes. Mol Cell 8, 1105–1115.

Deng, S.K., Gibb, B., de Almeida, M.J., Greene, E.C., and Symington, L.S. (2014). RPA antagonizes microhomology-mediated repair of DNA double-strand breaks. Nat Struct Mol Biol 21, 405–412.

Deng, S.K., Yin, Y., Petes, T.D., and Symington, L.S. (2015). Mre11-Sae2 and RPA Collaborate to Prevent Palindromic Gene Amplification. Mol Cell 60, 500–508.

Desai-Mehta, A., Cerosaletti, K.M., and Concannon, P. (2001). Distinct functional domains of nibrin mediate Mre11 binding, focus formation, and nuclear localization. Mol Cell Biol 21, 2184–2191.

Deshpande, R.A., Williams, G.J., Limbo, O., Williams, R.S., Kuhnlein, J., Lee, J.H., Classen, S., Guenther, G., Russell, P., Tainer, J.A., et al. (2014). ATP-driven Rad50 conformations regulate DNA tethering, end resection, and ATM checkpoint signaling. The EMBO journal 33, 482–500.

Dungrawala, H., Rose, K.L., Bhat, K.P., Mohni, K.N., Glick, G.G., Couch, F.B., and Cortez, D. (2015). The Replication Checkpoint Prevents Two Types of Fork Collapse without Regulating Replisome Stability. Mol Cell 59, 998–1010.

Falck, J., Coates, J., and Jackson, S.P. (2005). Conserved modes of recruitment of ATM, ATR and DNA-PKcs to sites of DNA damage. Nature 434, 605–611.

Ferrari, M., Dibitetto, D., De Gregorio, G., Eapen, V.V., Rawal, C.C., Lazzaro, F., Tsabar, M., Marini, F., Haber, J.E., and Pellicioli, A. (2015). Functional interplay between the 53BP1-ortholog Rad9 and the Mre11 complex regulates resection, end-tethering and repair of a double-strand break. PLoS Genet 11, e1004928.

Gnugge, R., Liphardt, T., and Rudolf, F. (2016). A shuttle vector series for precise genetic engineering of Saccharomyces cerevisiae. Yeast 33, 83–98.

Gobbini, E., Cassani, C., Villa, M., Bonetti, D., and Longhese, M.P. (2016). Functions and regulation of the MRX complex at DNA double-strand breaks. Microb Cell 3, 329–337.

Gobbini, E., Villa, M., Gnugnoli, M., Menin, L., Clerici, M., and Longhese, M.P. (2015). Sae2 Function at DNA Double-Strand Breaks Is Bypassed by Dampening Tel1 or Rad53 Activity. PLoS Genet 11, e1005685.

Goedhart, J., von Stetten, D., Noirclerc-Savoye, M., Lelimousin, M., Joosen, L., Hink, M.A., van Weeren, L., Gadella, T.W., Jr., and Royant, A. (2012). Structure-guided evolution of cyan fluorescent proteins towards a quantum yield of 93%. Nat Commun 3, 751.

Heidinger-Pauli, J.M., Unal, E., Guacci, V., and Koshland, D. (2008). The kleisin subunit of cohesin dictates damage-induced cohesion. Mol Cell 31, 47–56.

Hélène Tourrière, J.S., Armelle Lengronne and Philippe Pasero (2017). Single-molecule Analysis of DNA Replication Dynamics in Budding Yeast and Human Cells by DNA Combing. Bio-protocol 7, 17.

Hohl, M., Kochanczyk, T., Tous, C., Aguilera, A., Krezel, A., and Petrini, J.H. (2015). Interdependence of the rad50 hook and globular domain functions. Mol Cell 57, 479–491.

Hohl, M., Kwon, Y., Galvan, S.M., Xue, X., Tous, C., Aguilera, A., Sung, P., and Petrini, J.H. (2011). The Rad50 coiled-coil domain is indispensable for Mre11 complex functions. Nat Struct Mol Biol 18, 1124–1131.

Ivanov, E.L., Korolev, V.G., and Fabre, F. (1992). XRS2, a DNA repair gene of Saccharomyces cerevisiae, is needed for meiotic recombination. Genetics 132, 651–664.

Kaye, J.A., Melo, J.A., Cheung, S.K., Vaze, M.B., Haber, J.E., and Toczyski, D.P. (2004). DNA breaks promote genomic instability by impeding proper chromosome segregation. Curr Biol 14, 2096–2106.

Kim, J.H., Grosbart, M., Anand, R., Wyman, C., Cejka, P., and Petrini, J.H. (2017). The Mre11-Nbs1 Interface Is Essential for Viability and Tumor Suppression. Cell Rep 18, 496–507.

Kim, J.S., Krasieva, T.B., LaMorte, V., Taylor, A.M., and Yokomori, K. (2002). Specific recruitment of human cohesin to laser-induced DNA damage. J Biol Chem 277, 45149–45153

Krogh, B.O., Llorente, B., Lam, A., and Symington, L.S. (2005). Mutations in Mre11 phosphoesterase motif I that impair Saccharomyces cerevisiae Mre11-Rad50-Xrs2 complex stability in addition to nuclease activity. Genetics 171, 1561–1570.

Lammens, K., Bemeleit, D.J., Mockel, C., Clausing, E., Schele, A., Hartung, S., Schiller, C.B., Lucas, M., Angermuller, C., Soding, J., et al. (2011). The Mre11:Rad50 structure shows an ATP-dependent molecular clamp in DNA double-strand break repair. Cell 145, 54–66.

Lee, J.H., and Paull, T.T. (2005). ATM activation by DNA double-strand breaks through the Mre11-Rad50-Nbs1 complex. Science 308, 551–554.

Lee, K., Zhang, Y., and Lee, S.E. (2008). Saccharomyces cerevisiae ATM orthologue suppresses break-induced chromosome translocations. Nature 454, 543–546.

Liang, J., Suhandynata, R.T., and Zhou, H. (2015). Phosphorylation of Sae2 Mediates Forkhead-associated (FHA) Domain-specific Interaction and Regulates Its DNA Repair Function. J Biol Chem 290, 10751–10763.

Lim, H.S., Kim, J.S., Park, Y.B., Gwon, G.H., and Cho, Y. (2011). Crystal structure of the Mre11-Rad50-ATPgammaS complex: understanding the interplay between Mre11 and Rad50. Genes Dev 25, 1091–1104.

Limbo, O., Yamada, Y., and Russell, P. (2018). Mre11-Rad50-dependent activity of ATM/Tel1 at DNA breaks and telomeres in the absence of Nbs1. Mol Biol Cell 29, 1389–1399.

Lisby, M., Mortensen, U.H., and Rothstein, R. (2003). Colocalization of multiple DNA double-strand breaks at a single Rad52 repair centre. Nat Cell Biol 5, 572–577.

Lloyd, J., Chapman, J.R., Clapperton, J.A., Haire, L.F., Hartsuiker, E., Li, J., Carr, A.M., Jackson, S.P., and Smerdon, S.J. (2009). A supramodular FHA/BRCT-repeat architecture mediates Nbs1 adaptor function in response to DNA damage. Cell 139, 100–111.

Lobachev, K., Vitriol, E., Stemple, J., Resnick, M.A., and Bloom, K. (2004). Chromosome fragmentation after induction of a double-strand break is an active process prevented by the RMX repair complex. Curr Biol 14, 2107–2112.

Malone, R.E., Ward, T., Lin, S., and Waring, J. (1990). The RAD50 gene, a member of the double strand break repair epistasis group, is not required for spontaneous mitotic recombination in yeast. Curr Genet 18, 111–116.

Matsuzaki, K., Shinohara, A., and Shinohara, M. (2008). Forkhead-associated domain of yeast Xrs2, a homolog of human Nbs1, promotes nonhomologous end joining through interaction with a ligase IV partner protein, Lif1. Genetics 179, 213–225.

Mirzoeva, O.K., and Petrini, J.H. (2003). DNA replication-dependent nuclear dynamics of the Mre11 complex. Mol Cancer Res 1, 207–218.

Mockel, C., Lammens, K., Schele, A., and Hopfner, K.P. (2012). ATP driven structural changes of the bacterial Mre11:Rad50 catalytic head complex. Nucleic acids research 40, 914–927.

Morales, M., Theunissen, J.W., Kim, C.F., Kitagawa, R., Kastan, M.B., and Petrini, J.H. (2005). The Rad50S allele promotes ATM-dependent DNA damage responses and suppresses ATM deficiency: implications for the Mre11 complex as a DNA damage sensor. Genes Dev 19, 3043–3054.

Mozlin, A.M., Fung, C.W., and Symington, L.S. (2008). Role of the Saccharomyces cerevisiae Rad51 paralogs in sister chromatid recombination. Genetics 178, 113–126.

Myung, K., Chen, C., and Kolodner, R.D. (2001). Multiple pathways cooperate in the suppression of genome instability in Saccharomyces cerevisiae. Nature 411, 1073–1076.

Nakada, D., Matsumoto, K., and Sugimoto, K. (2003). ATM-related Tel1 associates with double-strand breaks through an Xrs2-dependent mechanism. Genes Dev 17, 1957–1962.

Oh, J., Al-Zain, A., Cannavo, E., Cejka, P., and Symington, L.S. (2016). Xrs2 Dependent and Independent Functions of the Mre11-Rad50 Complex. Mol Cell 64, 405–415.

Palmbos, P.L., Wu, D., Daley, J.M., and Wilson, T.E. (2008). Recruitment of Saccharomyces cerevisiae Dnl4-Lif1 complex to a double-strand break requires interactions with Yku80 and the Xrs2 FHA domain. Genetics 180, 1809–1819.

Park, Y.B., Chae, J., Kim, Y.C., and Cho, Y. (2011). Crystal structure of human Mre11: understanding tumorigenic mutations. Structure 19, 1591–1602.

Putnam, C.D., and Kolodner, R.D. (2010). Determination of gross chromosomal rearrangement rates. Cold Spring Harb Protoc 2010, pdb prot5492.

Ritchie, K.B., and Petes, T.D. (2000). The Mre11p/Rad50p/Xrs2p complex and the Tel1p function in a single pathway for telomere maintenance in yeast. Genetics 155, 475–479.

Roset, R., Inagaki, A., Hohl, M., Brenet, F., Lafrance-Vanasse, J., Lange, J., Scandura, J.M., Tainer, J.A., Keeney, S., and Petrini, J.H. (2014). The Rad50 hook domain regulates DNA damage signaling and tumorigenesis. Genes Dev 28, 451–462.

Ruff, P., Donnianni, R.A., Glancy, E., Oh, J., and Symington, L.S. (2016). RPA Stabilization of Single-Stranded DNA Is Critical for Break-Induced Replication. Cell Rep 17, 3359–3368.

Schiller, C.B., Lammens, K., Guerini, I., Coordes, B., Feldmann, H., Schlauderer, F., Mockel, C., Schele, A., Strasser, K., Jackson, S.P., et al. (2012). Structure of Mre11-Nbs1 complex yields insights into ataxia-telangiectasia-like disease mutations and DNA damage signaling. Nat Struct Mol Biol 19, 693–700.

Seeber, A., Hegnauer, A.M., Hustedt, N., Deshpande, I., Poli, J., Eglinger, J., Pasero, P., Gut, H., Shinohara, M., Hopfner, K.P., et al. (2016). RPA Mediates Recruitment of MRX to Forks and Double-Strand Breaks to Hold Sister Chromatids Together. Mol Cell 64, 951–966.

Shiloh, Y., and Ziv, Y. (2013). The ATM protein kinase: regulating the cellular response to genotoxic stress, and more. Nature reviews Molecular cell biology 14, 197–210.

Sirbu, B.M., Couch, F.B., Feigerle, J.T., Bhaskara, S., Hiebert, S.W., and Cortez, D. (2011). Analysis of protein dynamics at active, stalled, and collapsed replication forks. Genes Dev 25, 1320–1327.

Sjogren, C., and Nasmyth, K. (2001). Sister chromatid cohesion is required for postreplicative double-strand break repair in Saccharomyces cerevisiae. Current biology : CB 11, 991–995.

Smith, S., Gupta, A., Kolodner, R.D., and Myung, K. (2005). Suppression of gross chromosomal rearrangements by the multiple functions of the Mre11-Rad50-Xrs2 complex in Saccharomyces cerevisiae. DNA Repair (Amst) 4, 606–617.

Stewart, G.S., Maser, R.S., Stankovic, T., Bressan, D.A., Kaplan, M.I., Jaspers, N.G., Raams, A., Byrd, P.J., Petrini, J.H., and Taylor, A.M. (1999). The DNA double-strand break repair gene hMRE11 is mutated in individuals with an ataxia-telangiectasia-like disorder. Cell 99, 577–587.

Stracker, T.H., Morales, M., Couto, S.S., Hussein, H., and Petrini, J.H. (2007). The carboxy terminus of NBS1 is required for induction of apoptosis by the MRE11 complex. Nature 447, 218–221.

Stracker, T.H., and Petrini, J.H. (2011). The MRE11 complex: starting from the ends. Nature reviews Molecular cell biology 12, 90–103.

Strom, L., Karlsson, C., Lindroos, H.B., Wedahl, S., Katou, Y., Shirahige, K., and Sjogren, C. (2007). Postreplicative formation of cohesion is required for repair and induced by a single DNA break. Science 317, 242–245.

Strom, L., and Sjogren, C. (2007). Chromosome segregation and double-strand break repair - a complex connection. Curr Opin Cell Biol 19, 344–349.

Symington, L.S., Rothstein, R., and Lisby, M. (2014). Mechanisms and regulation of mitotic recombination in Saccharomyces cerevisiae. Genetics 198, 795–835.

Tittel-Elmer, M., Alabert, C., Pasero, P., and Cobb, J.A. (2009). The MRX complex stabilizes the replisome independently of the S phase checkpoint during replication stress. The EMBO journal 28, 1142–1156.

Tittel-Elmer, M., Lengronne, A., Davidson, M.B., Bacal, J., Francois, P., Hohl, M., Petrini, J.H.J., Pasero, P., and Cobb, J.A. (2012). Cohesin association to replication sites depends on rad50 and promotes fork restart. Mol Cell 48, 98–108.

Tsukamoto, Y., Mitsuoka, C., Terasawa, M., Ogawa, H., and Ogawa, T. (2005). Xrs2p regulates Mre11p translocation to the nucleus and plays a role in telomere elongation and meiotic recombination. Mol Biol Cell 16, 597–608.

Unal, E., Arbel-Eden, A., Sattler, U., Shroff, R., Lichten, M., Haber, J.E., and Koshland, D. (2004). DNA damage response pathway uses histone modification to assemble a double-strand break-specific cohesin domain. Mol Cell 16, 991–1002.

Unal, E., Heidinger-Pauli, J.M., and Koshland, D. (2007). DNA double-strand breaks trigger genome-wide sister-chromatid cohesion through Eco1 (Ctf7). Science 317, 245–248.

Waltes, R., Kalb, R., Gatei, M., Kijas, A.W., Stumm, M., Sobeck, A., Wieland, B., Varon, R., Lerenthal, Y., Lavin, M.F., et al. (2009). Human RAD50 deficiency in a Nijmegen breakage syndrome-like disorder. Am J Hum Genet 84, 605–616.

Williams, B.R., Mirzoeva, O.K., Morgan, W.F., Lin, J., Dunnick, W., and Petrini, J.H. (2002). A murine model of Nijmegen breakage syndrome. Curr Biol 12, 648–653.

Williams, R.S., Dodson, G.E., Limbo, O., Yamada, Y., Williams, J.S., Guenther, G., Classen, S., Glover, J.N., Iwasaki, H., Russell, P., et al. (2009). Nbs1 flexibly tethers Ctp1 and Mre11-Rad50 to coordinate DNA double-strand break processing and repair. Cell 139, 87–99.

Wiltzius, J.J., Hohl, M., Fleming, J.C., and Petrini, J.H. (2005). The Rad50 hook domain is a critical determinant of Mre11 complex functions. Nat Struct Mol Biol 12, 403–407.

You, Z., Chahwan, C., Bailis, J., Hunter, T., and Russell, P. (2005). ATM activation and its recruitment to damaged DNA require binding to the C terminus of Nbs1. Mol Cell Biol 25, 5363–5379.

## References

Mimitou, E.P., and Symington, L.S. (2008). Sae2, Exo1 and Sgs1 collaborate in DNA double-strand break processing. Nature 455, 770–774.

Yu, C., Gan, H., Han, J., Zhou, Z.X., Jia, S., Chabes, A., Farrugia, G., Ordog, T., and Zhang, Z. (2014). Strand-specific analysis shows protein binding at replication forks and PCNA unloading from lagging strands when forks stall. Mol Cell 56, 551–563.

